# 3D nanoscale architecture of the respiratory epithelium reveals motile cilia-rootlets-mitochondria axis of communication

**DOI:** 10.1101/2024.09.08.611854

**Authors:** Aaran Vijayakumaran, Christopher Godbehere, Analle Abuammar, Sophia Y. Breusegem, Leah R. Hurst, Nobuhiro Morone, Jaime Llodra, Melis T. Dalbay, Niaj M. Tanvir, K. MacLellan-Gibson, Chris O’Callaghan, Esben Lorentzen, CellMap Project Team, FIB-SEM Technology, Andrew J. Murray, Kedar Narayan, Vito Mennella

**Affiliations:** MRC Toxicology Unit, School of Biological Sciences, University of Cambridge; Cambridge, CB2 1QR, United Kingdom; Department of Molecular Biology and Genetics, Aarhus University; Aarhus, 8000, Denmark, EU; Babraham Hall House, Babraham; Cambridge, CB22 3AT, United Kingdom; Great Ormond Street Institute of Child Health, and GOSH BRC, University College London; London, WC1N 1EH, UK; Janelia Research Campus, Howard Hughes Medical Institute, Ashburn, Virginia, United States; Department of Physiology, Development and Neuroscience, University of Cambridge; Cambridge CB2 3EL, UK; Center for Molecular Microscopy, Center for Cancer Research, National Cancer Institute, National Institutes of Health; Bethesda, MD 20892, USA; Department of Pathology, University of Cambridge; Cambridge, CB2 1QP, United Kingdom

**Author notes:** These authors contributed equally to this work (Aaran Vijayakumaran, Christopher Godbehere).

## Abstract

A major frontier in single cell biology is decoding how transcriptional states result in cellular-level architectural changes, ultimately driving function. A remarkable example of this cellular remodelling program is the differentiation of airway stem cells into the human respiratory multiciliated epithelium, a tissue barrier protecting against bacteria, viruses and particulate matter. Here, we present the first isotropic three-dimensional map of the airway epithelium at the nanometre scale unveiling the coordinated changes in cellular organisation, organelle topology and contacts, occurring during multiciliogenesis. This analysis led us to discover a cellular mechanism of communication whereby motile cilia relay mechanical information to mitochondria through striated cytoskeletal fibres, the rootlets, to promote effective ciliary motility and ATP generation. Altogether, this study integrates nanometre-scale structural, functional and dynamic insights to elucidate fundamental mechanisms responsible for airway defence.

## Introduction

The human airway pseudostratified epithelium serves as the first line of defence against bacteria, viruses, and environmental pollutants inhaled through breathing (*1*). Recent studies employing single cell RNA sequencing (scRNAseq) have elucidated the transcriptional changes that occur as airway basal stem cells differentiate into a functional airway epithelium, establishing the existence of discrete cellular states (*2*). During this differentiation process, airway basal stem cells transition from an ellipsoid to an elongated shape along an apical-basolateral polarity axis, ultimately giving rise to six main cell types. However, how these transcriptional changes result in functionally relevant alterations of the three-dimensional structure and organisation of the epithelium remain unclear.

Here we focus on the process of transformation of a basal stem cell into the airway multiciliated cell, a crucial player in mucociliary clearance, the protective mechanism of the respiratory epithelium driven by beating motile cilia (*3*). Dysfunction of airway multiciliated cells contributes to the pathogenesis of lung diseases such as Primary Ciliary Dyskinesia (PCD) (*4*), asthma (*5*,*6*) and chronic obstructive pulmonary disease (COPD) (*7*,*8*).

During the multiciliogenesis trajectory, airway basal stem cells first progress into deuterosomal cells (*9*), an intermediate state where hundreds of basal bodies are generated in the cytoplasm, transported towards the plasma membrane and docked in patches (*10*). To form functional multiciliated cells, basal bodies then extend outward axonemes, membrane-bound microtubule-based scaffolds required for beating, forming mature motile cilia (*3*,*11*). Concomitantly, basal bodies extend inward toward the cytosol rootlets, cytoskeletal fibres whose organisation and function in human airway cells remain unclear. Although the main stages of the multiciliogenesis program have been identified, how airway cells reconfigure their internal organelles and cytoskeletal organisation to drive this process forward remains unknown.

Volume Electron Microscopy (vEM) technologies provide an opportunity to transform the view of cellular differentiation trajectories, providing detailed nanometre resolution information *in situ* within the tissue milieu, aiding in the development of mechanistic hypotheses in a non-reductionist manner (*12*). In this study, we employed an unbiased 3D volume segmentation approach to uncover how organelle and cytoskeletal elements work as an ensemble to determine the mechanisms responsible for multiciliated cell formation and its essential mucociliary clearance function.

## Results

### Focused ion beam-scanning electron microscopy (FIB-SEM) and 3D segmentation reveals multiscale changes during multiciliogenesis

To systematically analyse cellular changes occurring during the formation of an airway multiciliated cell, we developed a 3D volume segmentation, cell extraction, and visualisation pipeline for analysis of the human airway pseudostratified epithelium, a tissue reconstituted *in vitro* through an air-liquid interface culture system (*13*) (fig.1a, S1a). Volume Electron Microscopy (vEM) isotropic datasets, acquired by FIB-SEM, were reconstructed in 3D (fig.1b, movie 1,2) through the Python package and Napari Plugin, Empanada (*14*). This tool, leveraging a panoptic deep lab model pretrained with 1.5 million EM images, enabled the creation of airway epithelium-specific image models, recognising organelles and structures of interest based on a subset of manually segmented images (fig.1c) (*15*). Discrete multiciliogenesis cellular states were identified based on the presence of state-specific organelles (e.g., deuterosome, motile cilia) or position within the epithelium (fig.1d-g, S1b, c) (*3*,*11*,*16*). Upon 3D inference of the entire volume, individual cells were extracted using Python script using a membrane boundary mask.

3D nanoscale architecture analysis of the cell states along the multiciliogenesis trajectory revealed extensive rearrangements in cellular content, organelle topology and interactions, and organisation of supramolecular structures (fig.1d, S2a, b). Notable alterations pertain to both the plasma membrane (fig. S3,4) and perinuclear localisation of organelles (fig.2a,b, S5). In basal stem cells, the plasma membrane exhibits a predominantly ellipsoid organisation. During differentiation however, the apical membrane undergoes a transition from a concave architecture in deuterosomal cells (figs S3a-c), marked by frequent recycling endosomes fusing with the plasma membrane (fig. S4a-c), to a convex dome-like architecture in multiciliated cells, suggesting membrane deposition and extension over differentiation (fig. S3a-c).

The nuclear architecture also undergoes substantial changes during differentiation, becoming progressively more elongated in the apical-basolateral axis direction, while the nuclear membrane features extensive nuclear invaginations in all stages, showing wrinkling characteristic of low-tension conditions (*17*). Interestingly, in basal stem cells the invaginations are filled with endoplasmic reticulum and mitochondria, indicative of mitochondria-nuclear communication (*18*) via local metabolic activity and translation (*17*). This differs from differentiated cells, whose nuclei are surrounded by numerous lipid droplets (fig. 2b fig. S5a), consistent with the requirement of multiciliated cells for rapid metabolic rewiring toward fatty acid beta oxidation during repair (*19*).

**Figure 1.**
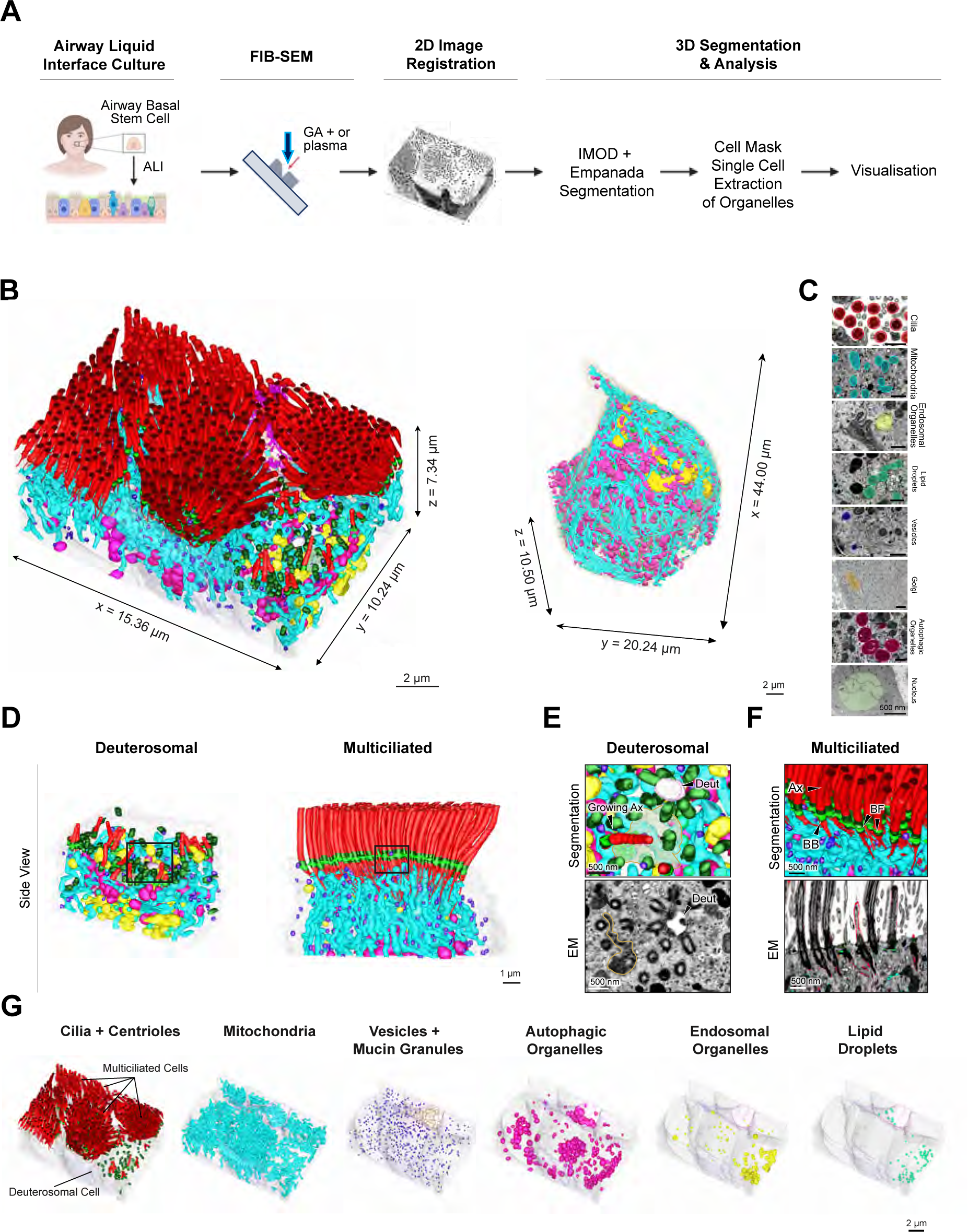
(A) Workflow for FIB-SEM 3D reconstructions of the human airway epithelium. **(B)** 3D volume segmentation of the human nasal airway epithelium containing deuterosomal and differentiated multiciliated (MCC) cells (left), and airway basal stem progenitor cell (right). **(C)** Examples of organelles identification on 2D EM micrographs using segmentation masks. **(D)** 3D volume segmentations of deuterosomal and MCC cell apical regions shown in side view **(E)** Top. High-magnification of inset from (D) showing 3D reconstruction of deuterosomal-mediated centriolar amplification. Arrowheads point to the deuterosome (Deut), and a growing axoneme (Ax). Bottom. EM image corresponding to top panel overlaid with segmentation contours. **(F)** Top. High-magnification of inset from (D) from MCC volume. Arrowheads point to the fully formed axonemes (Ax), basal bodies (BB) and aligned basal feet (BF). Bottom. EM image corresponding to top panel overlaid with segmentation contours. **(G)** Organelle breakdown of the 3D FIB-SEM reconstruction from (B).

**Figure 2:**
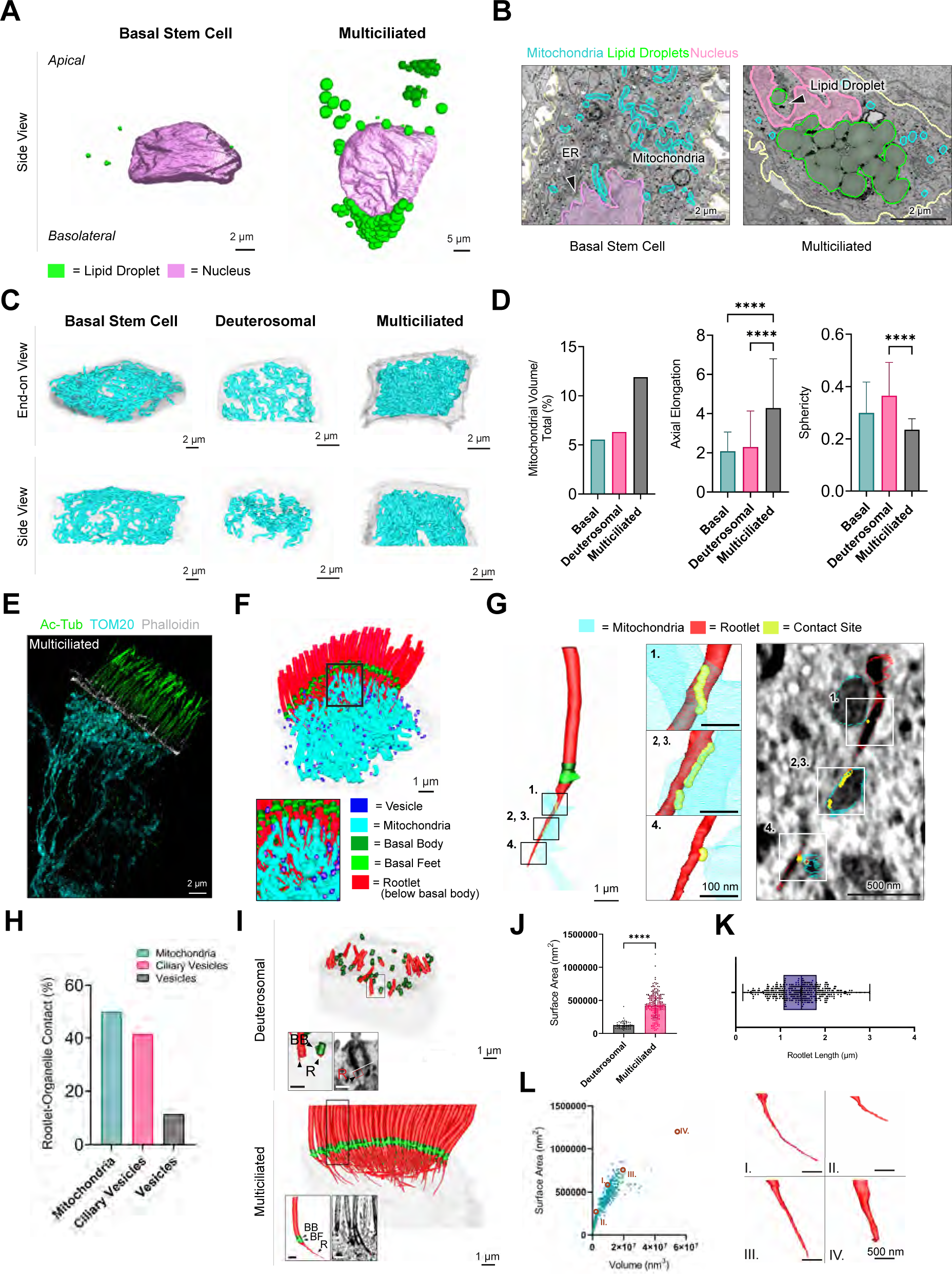
(A) 3D volume segmentation of nucleus (pink) and lipid droplets (green) in airway basal stem cell and MCC. (B) 2D FIB-SEM slices shown in end-on view highlighting differences in organelle content around the nuclear envelope and within nuclear invaginations between airway basal stem cells and MCCs. (C) 3D volume segmentation of plasma membrane (grey) and mitochondria (blue) in the apical regions of a deuterosomal cell and MCC, and in an airway basal stem cell. (D) Quantitative comparison of mitochondria volume and shape features in differentiated MCC (n=45), deuterosomal cell (n=42) and basal stem cell (n=40). Data were analysed with Kruskal-Wallis ANOVA and Dunn’s multiple comparisons test. **** p < 0.0001. (E) 3D-SIM volume maximum intensity projection of human MCC labelled with anti-TOM20 (cyan), anti-Ac-tubulin (Ac-Tub; green), and Phalloidin (grey) antibodies. (F,G) 3D volume segmentation of mitochondria-rootlets contacts. (H) Quantitative analysis of frequency of rootlet-organelle contacts (n=587 contacts in n=4 differentiated MCCs). (I) 3D volume segmentation of axonemes (red, above the plasma membrane), basal bodies (green) and ciliary rootlets (red, below the plasma membrane). The inset shows a high magnification view of an individual cilium and associated rootlet, overlaid with segmentation contours from an EM slice. Scale bar of segmentation inset: 500nm. Scale bar of EM inset: 200 nm. Notations: BB (basal body), BF (basal foot), R (rootlet). (J) Comparison of surface area between Deuterosomal (n = 54 rootlets) and Multiciliated Cell (n = 223 rootlets). Data were analysed with the Mann-Whitney u-test. p<0.0001. (K) Distribution of the rootlets’ length, n=411. (J) Surface area-to-volume relationship for 411 ciliary rootlets obtained from n = 4 differentiated MCCs with examples of the specific ciliary rootlets (I-IV) showing diverse morphological characteristics.

3D volume segmentation allowed further observation of several structural changes linked to basal body maturation occurring during deuterosomal to multiciliated cell transition. In deuterosomal cells, migrating centrioles and docked basal bodies extend a single, narrow sub-distal appendage, which subsequently remodels into a wide cone-structured basal foot in fully differentiated cells (fig. S6a,b). This observation clarifies the timing of structural reorganisation of the subdistal appendage to the basal foot, a supramolecular assembly critical for motile cilia coordination that links basal bodies through microtubule cables (*20*,*21*). Interestingly, in deuterosomal cells, migrating centrioles and docked basal bodies lack the distinct ciliary vesicle typical of primary cilia required for its extension (fig. S4d) (*22*).

Moreover, basal bodies in multiciliated cells present an electron dense elongated structure, which is positioned toward the distal end of the basal body between the transition fibres and basal feet in the longitudinal direction, and in the centre of the basal body in the lateral direction (fig. S7a, b). This structure is fully formed in extended axonemes and is positioned within the motile cilia lumen suggesting that is functionally linked to motile cilia beating and is made of a protein complex distinct from inner helical scaffold proteins associated with axonemal microtubules such as POC1 and POC5 (*23*) (fig. S7c-e).

### The mitochondrial network and contacts remodel upon multiciliogenesis

A remarkable difference between multiciliogenesis cell states lies in the density, distribution, and shape of the mitochondrial network. Quantitative analysis of volumetric data (fig. 2c, d, S8a) and super-resolution imaging (fig.2e, S8b) revealed that mitochondria exhibit highly elongated organisation along the apical-basolateral axis in multiciliated cells, which contrasts with the lateral distribution observed in deuterosomal cells and the homogenous distribution in basal stem cells (fig. 2c,d, S8b). Notably, the mitochondrial network becomes densely packed near the apical membrane in multiciliated cells, consistent with the role of mitochondria in generating the ATP required to propel motile cilia (fig. 2e) (*24*).

Unexpectedly, at the apical surface of multiciliated cells, the mitochondrial network forms a highly interconnected network with the rootlets, striated cytoskeletal fibres extending from the basal bodies of motile cilia (fig. 2f,g S8c,d). The mitochondria make either a single contact or wrap around rootlets along their length (fig. 2g, S8c-d), with more than 50% of the rootlets making contact with mitochondria and less frequently with ciliary vesicles (2h, S8c-e). The mitochondria-rootlets interactions are reminiscent of the contacts observed in mouse photoreceptor cilia and marine invertebrates (*25*,*26*), suggesting functional interplay between the organelle and cytoskeletal fibre.

In multiciliated cells, the rootlets form fibres of varying lengths (1.45 ± 0.5 µm), thickness, volume, and surface area, which contrasts with the stubs emanating from basal bodies in deuterosomal cells (fig. 2i-l, S8f-h, movie 3). Despite their length and curvature (fig. 3a), the rootlets do not intertwine with each other, as observed in cycling cells (*27*), do not contact the nucleus as in other cell types (*28*) and extend mainly longitudinally. This organisation differs from the striated fibres, analogous structures in evolutionarily more ancient organisms, which extend laterally right below the apical surface and connect basal bodies together (*29-31*). Analysis of individual rootlet origin and endpoint showed a highly ordered 3D spatial organisation, with fibres converging toward the posterior-end of the cell relative to the effective stroke, as indicated by the tip of the basal feet, which point toward the direction of motile cilia beating (fig. 3b, S9) (*10*,*32*). This suggests that the intracellular architecture of rootlets is functionally linked to the mechanism of ciliary motility occurring extracellularly.

**Figure 3.**
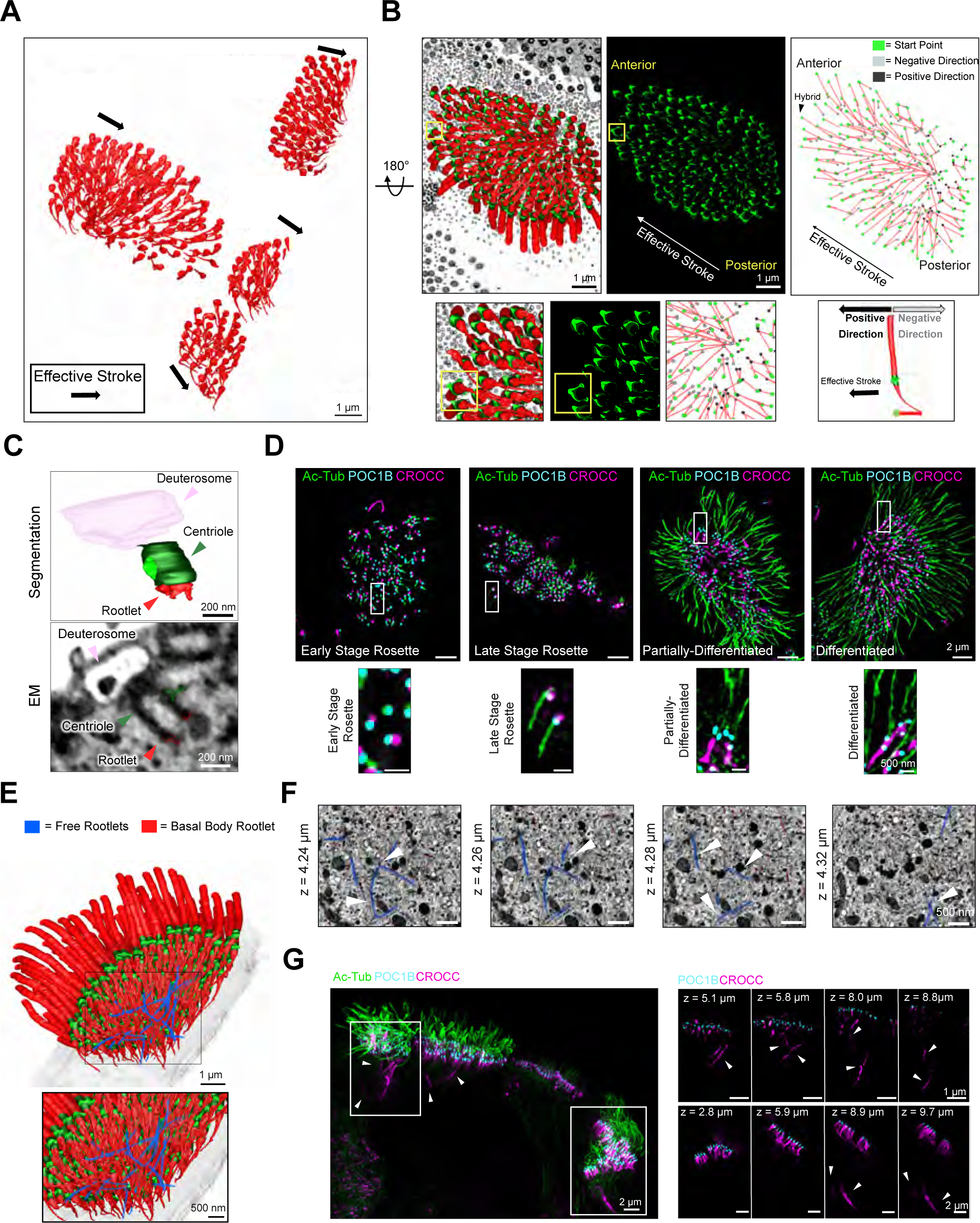
**(A)** 3D volume segmentation of ciliary rootlets in MCCs. **(B)** 3D volume segmentation of a differentiated MCC viewed from the bottom after 180 degrees rotation, showing ciliary rootlets (left panel) and basal feet alignment (middle panel) to determine effective stroke direction. In the right panel rootlets origins, endpoint and curvature direction are labelled on the 3D volume segmentation (n = 223). Note the convergence of rootlets toward the posterior end (away from the power stroke direction) and the hybrid cilium at the anterior end of the cell, which is preferentially found toward the direction of the power stroke. **(C)** 3D volume reconstruction of a migrating centriole with a ciliary rootlet stub extending from a deuterosomal cell. Below is the corresponding EM orthoslice overlaid with segmentation contours. **(D)** Top. Maximum intensity projection from 3D-SIM^2^ volume of human MCCs at different stages of differentiation labelled with POC1B (cyan) Acetylated-tubulin (green) and CROCC (magenta). Bottom. High magnifications of white insets. **(E)** Top. 3D volume segmentation of a differentiated MCC highlighting plasma membrane (grey), basal bodies (green) motile cilia (red above the basal bodies) and rootlets (red, below the basal bodies) and a subpopulation of free cytoplasmic rootlets (blue). Bottom. High magnification of white insets. **(F)** EM volume 2D slices with segmentation contours highlighting examples of free rootlets. **(G)** Left panel: maximum intensity projection of 3D-SIM^2^ micrographs of human MCCs, labelled with POC1B (cyan), Ac-Tub (green), and CROCC (magenta). Right: series of z-sections from insets in the left panel. Free rootlets are labelled with white arrowheads.

### Supramolecular architecture and assembly of rootlets across resolution scales

The unexpected cellular organisation and interactions of the rootlets prompted us to further investigate their assembly, nanoscale organisation, and function during multiciliogenesis.

To examine when rootlets form and elongate, airway multiciliated cells at different differentiation stages were labelled with antibodies anti-CROCC/Rootletin, the main constituent of rootlets, (*33*) and imaged via lattice 3D-Structured illumination microscopy (3D-SIM), a super-resolution microscopy featuring multicolour imaging below the diffraction limit (*34*). Consistent with 3D segmentation analysis (fig. 3c), rootlets emerge as short stubs during basal body duplication and start elongating after basal body docking and partial axonemal extension (fig. 3d). Interestingly, vEM and super-resolution microscopy experiments both revealed a subset of rootlets that are not tethered to basal bodies, which we termed ‘free rootlets’ (fig. 3e-g, S10a, b, movie 4,5). These rootlets are positioned subapically and partly colocalise with PCM1, a protein previously implicated in phase separation in ependymal multiciliated cells (fig. S10c) (*35*). Untethered free rootlets are also observed in airway multiciliated cells lacking basal bodies due to CCNO loss of function, a transcription factor critical for basal body duplication (fig. S10d-f) (*36*,*37*). These observations support the view that rootlets are polymers capable of self-assembly within cells independently of basal bodies, with the latter structures being responsible for anchoring and directing rootlet extension.

Next, we examined how rootlets are molecularly linked to basal bodies. 3D segmentation of FIB-SEM data shows that rootlet-like fibrils originate from distinct regions of the outer diameter of the basal body’s proximal end, rapidly coalescing into a single bundle tapering into the cytoplasm (fig. S11a). By using High-Pressure Freezing and Freeze Substitution Transmission Electron Microscopy (HPF-FS-TEM) we however revealed a more complex network characterised by fibrils emanating from the basal body central region (fig. 4a inset 1) and outer diameter (fig. 4a inset 2), indicating that unidentified proteins may contribute to its structure. To examine the organisation of this linker structure, a nanoscale mapping approach was employed to first determine the relative positions of proteins previously implicated in bridging basal bodies and rootlets using the distal basal body marker POC1B as a reference (S11b, c) (*33*,*38–43*). 3D-SIM^2^ imaging shows that coiled-coil protein CEP250 resides between the proximal end of the basal body, labelled by CEP135, and the rootlets, labelled by CROCC (fig. 4b, c, S11d). CEP250 molecules exhibit an elongated molecular architecture, with their C-termini closer to the basal body (316 nm ± 20.2 nm) and N-termini (458.5 nm ± 34.5 nm) in proximity to the C-terminus of CROCC (510.9 nm ± 67.6 nm) (fig. 4b, c, S11d), in agreement with CEP250 being part of the molecular linker between the basal body and the rootlets (*43*). Conversely, CROCC binding partner LRRC45 is distributed uniformly throughout the linker region in addition to its transitional fibre localization (fig. 4d) (*44*), while CEP44 is located much closer to the basal body (186.1 nm ± 20.8 nm) than CEP250 and CEP135 (284 nm ± 33.9 nm) (fig. 4b, c, S11d, Table S2).

**Figure 4.**
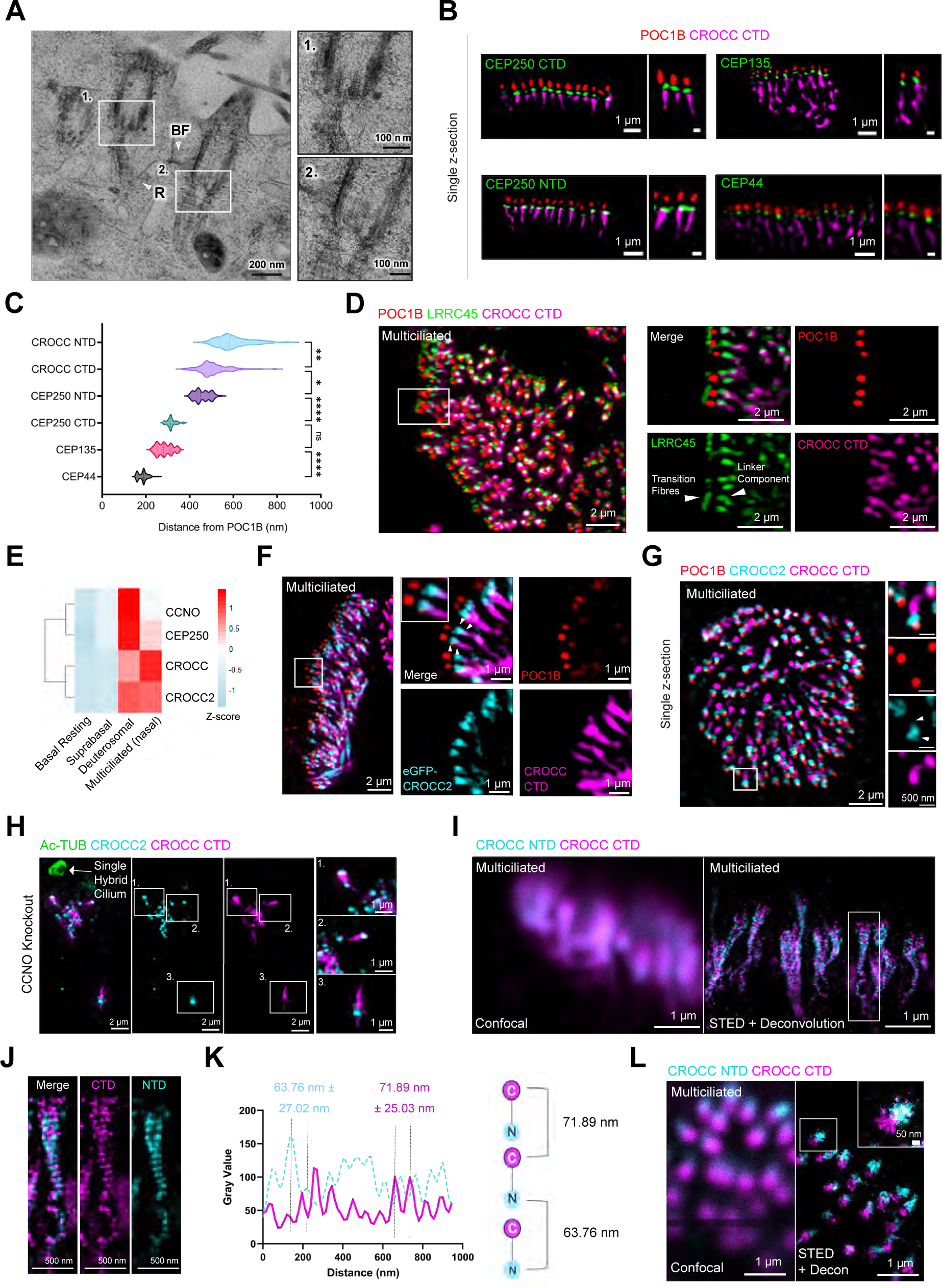
(A) TEM micrographs of HPF-FS MCCs, showing docked centrioles and rootlets (arrowheads) emanating from the perimeter and the centre of the proximal end of the basal body (BF, basal foot; R, Rootlet.) (B) Single z-sections from 3D-SIM2 volumes of MCCs labelled with POC1B as a reference marker, CROCC and different proteins previously suggested to link basal bodies and rootlets. Scale bar of the high magnification insets: 250 nm (C) Distance measurements from centriolar marker POC1B of proteins suggested to link basal bodies and rootlets (n = 97-129 centrioles, n = 10-14 cells) (See Table S2). Data were analysed with Kruskal-Wallis ANOVA and Dunn’s multiple comparisons test. * p = 0.0464, ** p = 0.0022, **** p < 0.0001. (D) 3D-SIM volume maximum intensity projection of a differentiated MCC labelled with anti-LRRC45 (green), POC1B (red) and CROCC CTD (magenta) antibodies. Inset in white box shows high magnification of rootlets on their side emphasising localisation of LRRC45 at both the transition fibres and proximal end of the centriole where it forms filaments overlapping with CROCC CTD (arrowheads). (E) Differences in expression (log2-fold changes) of CCNO, CEP250, CROCC2 and CROCC during human airway multiciliated differentiation. (F & G) 3D-SIM volume maximum intensity projection and single z-section of MCCs labelled with POC1B (red), eGFP-CROCC2 (cyan), antibody anti-CROCC2 (cyan), and CROCC CTD (magenta). Arrowheads point to the linkage between the rootlets and the basal bodies. (H) Maximum intensity projection of 3D-SIM volume from CCNO KO human airway cells labelled with Acetylated-alpha-tubulin (green), CROCC CTD (magenta), and CROCC2 antibody (cyan). Note high-magnification insets showing CROCC2 accumulation at one end of the cytoplasmic rootlets labelled by CROCC antibodies. (I) Confocal or 2D STED plus deconvolution imaging of CROCC NTD (cyan) and CTD (magenta) in a MCC imaged on side view. (J) High-magnification of image shown in (I) highlighting the periodic distribution of N- and C-termini of CROCC in a single ciliary rootlet. (K) Line intensity profile of the rootlets filament in (I) showing distances of CROCC N-N termini (63.8 nm ± 27.0 nm; n=355; n=21 rootlets) and C-C termini (71.9 nm ± 25.0 nm; n=314; n=21 rootlets). (L) 2D STED plus deconvolution imaging of CROCC N- and C-termini in a MCC in the top-down view. Note the lateral displacement between the two termini in the inset high magnification.

To identify new components of the basal body-rootlet linker region, we then examined CROCC2, a paralogue of CROCC and CEP250 (*45*). *CROCC2* encodes a protein of 1653 amino acids characterised by extensive coiled-coil domains, whose function and subcellular localisation are unknown (*46*). Analysis of scRNAseq data from Human Lung Cell Atlas shows that *CROCC2* transcript is highly expressed in multiciliated cells and follows a similar expression profile to CROCC and CEP250 (fig. 4e), therefore suggesting that it can be a *bona fide* component of rootlets. To test this hypothesis, we performed super-resolution imaging of multiciliated cells labelled with CROCC2 antibodies or expressing eGFP-CROCC2. These experiments showed that CROCC2 forms a structure linking the basal body to rootlets labelled by CROCC, both in migrating and docked basal bodies and when extending axonemes (fig. 4f,g, fig. S11e). CROCC2 forms a toroid structure located in proximity of the basal body with a short filament extending toward and partly overlapping with CROCC, suggesting a molecular interaction (fig. 4f,g white arrowheads). Remarkably, in cells lacking basal bodies due to CCNO KO, rootlet fibres assembling in the cytoplasm display CROCC2 accumulating at one end and less often in smaller dots along the fibre (fig. 4h), suggesting that CROCC2 labels the end of the rootlet independently of basal bodies, consistent with a role in establishing polarity to its structure.

As nanoscale mapping experiments suggested that CROCC molecules have an elongated organisation (fig. 4b), we next examined the position of its domains by antibody labelling and 2D STED microscopy plus deconvolution, which features higher in-plane resolution than lattice SIM (*34*). STED imaging of airway multiciliated rootlets imaged on the side revealed that CROCC assemblies form discrete repetitive units with N-termini and C-termini longitudinally separated by 63.4 nm and 71.5 nm, respectively, with C-termini located closer to the basal body, in agreement with previous studies in cycling cells and our SIM nanoscale mapping (fig. 4i-k) (*27*,*33*). Top-down imaging shows that the N- and C-termini of CROCC molecules display an offset in the axial direction (fig. 4l).

To link the nanometre scale *in situ* information with angstrom resolution structure, we performed structural prediction of CROCC multimers using AlphaFold v3.0. This analysis suggests that CROCC adopts the structure of a parallel homo-dimer along most of its 2000 residues (fig. S12a-c), which is consistent with a recent study (*47*), however coiled-coil helices between residues 600-1900 fold back yielding a rather compact 45 nm long helical bundle (fig. S12a). CROCC homo-dimer oligomerisation might lead to a more elongated structure; however, this possibility cannot be tested in AlphaFold due to constraints on input size. Altogether, the structure prediction is consistent with our *in situ* domain measurements by super-resolution microscopy supporting the notion that CROCC is not a fully extended (>250 nm) coiled coil protein.

### Rootlets counterbalance motile cilia forces in the cytoplasm and regulate mitochondria topology enhancing respiratory capacity

It has previously been shown that CROCC knockout (KO) mice exhibit increased infiltration of immune cells in the lung and are predisposed to respiratory infections (*48*), which suggested defects in mucociliary clearance. The architecture of rootlets, along with the timing of their extension coinciding with axonemal elongation, indicate that these cytoskeletal fibres might play a biomechanical role in ciliary motility, however the specific role of rootlets in airway multiciliated cells remains elusive.

To better understand if and how rootlets influence motile cilia coordination, airway multiciliated cells expressing eGFP-CROCC were generated using the PiggyBac integration system (fig. 5a). We reasoned that if rootlet fibres function in regulating ciliary beating forces, they might be dynamic structures responsive to ciliary motility. To test this possibility, eGFP-CROCC cells needed to be imaged with both high temporal resolution to follow cilia beating and spatial resolution to separate the individual rootlets. This analysis was made possible by using live super-resolution imaging by Lattice 3D-SIM with subsequent reconstruction by SIM^2^ algorithm (*49*). Our analysis revealed that rootlets undergo oscillatory movements with frequencies comparable to ciliary beating (9.4 Hz +/- 0.1) fig. 5b, movie 6). This oscillatory behaviour is coupled to ciliary motility, as inhibition of dynein motor function by Ciliobrevin D or addition of cross-linking agents blocked eGFP-CROCC movement.

**Figure 5.**
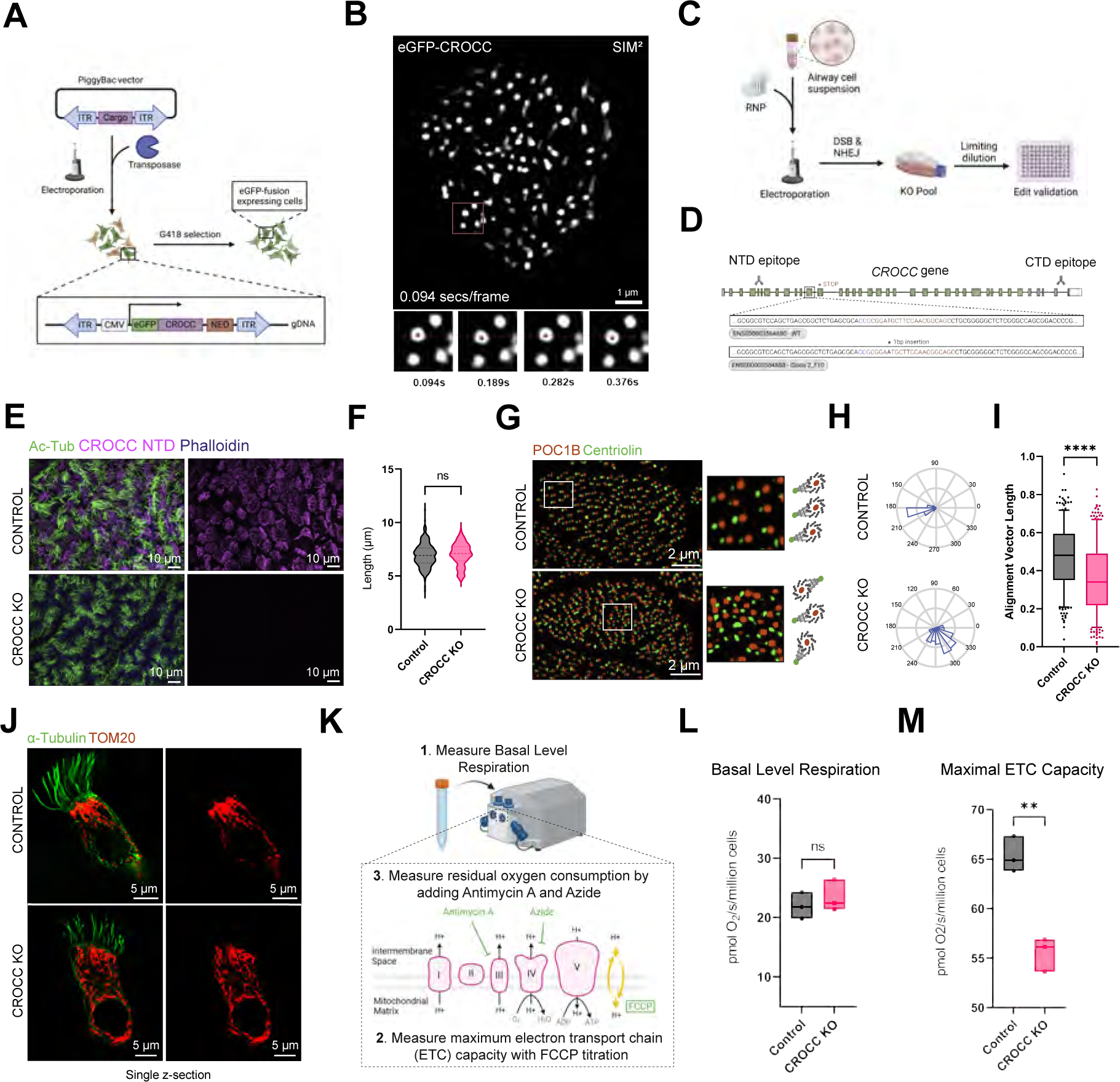
**(A)** Integration strategy for eGFP-CROCC expression in human airway cells. **(B)** eGFP-CROCC live-cell Lattice SIM imaging video reconstructed by SIM^2^ of a differentiated MCC imaged end-on. The bottom panels from the inset show a time series of a subset of eGFP-CROCC labelled rootlets undergoing oscillatory motion. The red dots mark the centroid of the rootlet, moving at an average speed of 9.8 Hz. **(C)** Workflow of the gene-targeting approach by CRISPR-Cas9. RNP (Ribonucleoprotein complex), DSB (Double-strand break), NHEJ (Non-homologous end-joining) **(D)** Location and sequence changes caused by CRISPR Cas9 edits on *CROCC.* Note that the 1 base pair insertion result in a premature stop codon in Exon 12 of the human *CROCC* gene **(E)** Maximum intensity projection from Airyscan fluorescence image stack obtained from control and CROCC knockout (KO) cells labelled with antibodies against anti-Ac-tubulin (green) and CROCC (magenta). **(F)** Cilia length distributions of fully differentiated CROCC KO cells (mean = 7.01 μm, n of cilia measured = 269, n of cells = 28) compared to control (mean = 6.95 μm, n of cilia measured = 267, n of cells = 29). Data were analysed with unpaired t-test. **(G)** Representative examples of maximum intensity projections from 3D SIM^2^ volumes of airway multiciliated cells from a control sample (Top) and CROCC KO (Bottom) labelled with basal body marker POC1B (red) and basal foot marker Centriolin (green) along with cartoon depiction of basal body/basal foot pairs alignment in control and CROCC KO cells. **(H)** Representative examples of rose plots from airway multiciliated cells from a control sample and CROCC KO cell. **(I)** Alignment vector length distribution of fully differentiated CROCC KO cells (mean = 0.36, n = 375 cells) compared to control (mean = 0.47, n = 372 cells). **** p < 0.0001. Data were analysed from n = 3 independent biological replicates with the Mann-Whitney u-test. **(J)** 3D-SIM volume maximum intensity projection of control and KO human airway cells labelled with Acetylated-alpha-tubulin (green) and TOM20 (red). **(K)** Strategy for high-resolution respirometry of fully differentiated airway epithelial cells to assess mitochondrial function between control and CROCC KO. **(L)** Basal respiration of fully differentiated CROCC KO cells (mean = 23.43) compared to control (mean = 21.96). **(M)** Maximal electron transport chain (ETC) capacity of fully differentiated CROCC KO cells (mean = 55.5) compared to control (mean = 65.2). ** p < 0.0024. Data were analysed with unpaired t-test from n = 3 independent biological replicates.

To directly test the hypothesis that rootlets are critical for coordinated beating of motile cilia, airway stem cells KO for *CROCC* were generated using CRISPR-Cas9, followed by selection of homozygous KO clones to enable robust phenotypic analysis (fig. 5c, d, S13). CROCC KO airway cells differentiate normally into multiciliated cells, thus suggesting that rootlets are not required for motile cilia assembly, differently from primary cilia (fig. 5e, S14) [47]. Moreover, CROCC KO cells exhibit similar motile cilia length relative to controls (fig 5f) and beating frequency (control 8.91 Hz +/- 1.6 n=3, 44 movies vs CROCC KO 8.89 Hz +/- 2.01 n=3,47 movies). Analysis of airway multiciliated cells differentiated for 55 days without rootlets also did not show overt differences in motile cilia stability, unlike primary cilia (*38*,*48*).

However, when motile cilia beating was inspected, CROCC KO cells appeared to show less coordinated beating than in healthy control cells. To quantitatively assess this phenotype, we analysed basal body/basal foot rotational polarity, a proxy for motile cilia coordination within a cell using automated MATLAB analysis on super-resolution micrographs of airway cells labelled with basal bodies and basal feet markers (*20*,*32*). This analysis revealed a robust reduction in alignment in CROCC KO cells (fig. 5g-i, Table S3), which is comparable to the misalignment caused by mutations in genes causing less severe forms of PCD, a disease characterised by immotile or dyskinetic cilia (*32*). Taken together, this shows that rootlets contribute to coordinated beating of motile cilia.

Since rootlets transduce the forces generated from motile cilia into the cytoplasm and form highly frequent and extensive contacts with mitochondrial membranes in multiciliated cells, it is tenable that rootlets might play a role in mitochondrial positioning within multiciliated cells and affect their metabolic activity. To test this possibility, we first analysed the organisation of the mitochondria by super-resolution microscopy. This analysis showed that mitochondria in CROCC KO cells do not concentrate apically as in control cells but are distributed more uniformly along the apical basolateral axis (fig.5j). Next, we employed high-resolution respirometry to measure mitochondrial respiratory capacity in the presence or absence of rootlets (fig. 5k). This analysis revealed that whilst basal level respiration was similar between control and CROCC KO cells, maximal electron transport chain capacity was significantly reduced (17.6%) in CROCC KO cells, indicating an impaired capacity to support oxidative phosphorylation in the absence of rootlets, resulting in lower ATP generation (fig. 5l,m, Table S4).

## Discussion

Here, we present the first 3D nanoscale map of the airway epithelium by vEM describing the cellular and molecular changes during the trajectory towards formation of a multiciliated cell, a key player in the lung defence mechanism against infectious agents and pollutants. *In situ* 3D cellular reconstruction by FIB-SEM transformed our view of this key process by unveiling multiple structural and topological changes in organelle and cytoskeleton interactions across differentiation, which will inform future studies in stem cell differentiation and comparisons of multiciliated epithelium organisation beyond the airway. To link cellular architectural changes to function, we focused on the analysis across scales of rootlets, one of the four main cytoskeletal fibres of the cell. While its main constituent — CROCC/Rootletin— and their striated appearance are conserved in multicellular organisms and mammals, their architecture and function remained poorly characterised in the airways.

Our data show that motile ciliary forces extend beyond the membrane region toward the cell’s interior, and rootlets function as cytoskeletal fibres transducing these forces to ensure beating coordination. It has been previously hypothesised that rootlets in lower organisms function as force-absorbing structures counteracting bending of basal bodies due to ciliary motility (*30*,*31*,*50*,*51*). In airway cells, however, rootlets do not connect basal bodies to each other as in striated fibres (*31*,*52*), a function played by basal feet in human multiciliated cells (*20*,*21*,*36*). Rather we found evidence that in human airway cells rootlets are active force-balancing structures rather than passive, immobile roots. Rootlets present a smaller range of motion relative to motile cilia as force is transmitted through a membrane-embedded basal body and they move in the intracellular milieu, which is characterised by different viscosity, density and molecular interactions relative to the extracellular environment.

The 3D reconstruction of human airway cells reveals a high-frequency organelle-cytoskeleton interaction involving rootlets and mitochondria. Rootlets align with and make multiple interactions with mitochondria, which are located apically for effective ATP delivery where it is most needed. This interaction is conserved across evolution as marine invertebrates such as lancelets present similar interaction with mitochondria, with cristae aligning with striations (*26*). Notably, our data provide evidence that rootlets are key for positioning mitochondria to the apical membrane and that the presence of rootlets around mitochondrial network supports greater oxidative phosphorylation capacity thereby linking for the first time the organelle-cytoskeletal fibre interaction to a key cell physiological function.

Recent CryoET studies have shown that rootlets form protrusions contacting membranes in mouse photoreceptor cells (*47*). This suggests the presence of CROCC domains and bridging protein complexes mediating interactions with membranous organelles including mitochondria, nuclei (*28*), golgi (*53*) and sacculous membrane (*48*). Nesprin-1 has been shown to mediate the interaction of rootlets with nuclei in photoreceptor cells (*28*). Altogether, this suggests the existence of protein complexes linking rootlets with mitochondria with a role in mechanotransduction (*54*), however, the molecular identity of the bridging protein complexes to mitochondria is not yet known.

Nanoscale imaging experiments show that rootlets are polar filaments capable of polymerising independently of basal bodies, yet their growth directionality is determined by their anchoring to the ciliary base. We observed rootlet elongation occurring after axonemal extension suggesting that their assembly is highly regulated in the “ciliary motility” molecular assembly line. The presence of free rootlets in proximity to existing rootlets in the apical region emanating from basal bodies suggests that rootlet elongation and assembly may be partly driven by cytoplasmic polymerisation of smaller rootlets and subsequent addition to the basal bodies, rather than relying solely on directional assembly. Consistent with this model, rootlets assemble filaments with bridging proteins such as CROCC2 at one end in the absence of basal bodies due to CCNO loss of function.

Our super-resolution mapping strongly supports a model where CROCC molecules have a fold back architecture and paralogues CEP250 and CROCC2 and LRRC45 form a supramolecular bridge linking basal bodies to the rootlets fibres made of CROCC. Consistent with this notion, structural modelling using AlphaFold suggest that CROCC2, CEP250 and CROCC can interact with each other as hetero-dimers through the formation of parallel coiled-coil structures, while LRRC45 interacts with CROCC through its leucine-rich N-terminal domain (fig. S15).

Altogether, this study provides a blueprint for future analyses aimed at defining tissue-level structural and functional maps in the respiratory tract. It represents the first step toward building a novel, extended atlas modelled upon single-cell RNA sequencing efforts, which will be integrated with volumetric electron microscopy data and other omics modalities, such as single-cell proteomics, to construct a comprehensive view of tissue structure and function.

## Acknowledgments

The authors gratefully acknowledge the Advanced Light Imaging, the Electron Microscopy and the Bioinformatics facility at the MRC Toxicology Unit for their support and assistance in this work. Dr Feride Oetztuerk-Winder for providing PiggyBac vector and for molecular biology help. The super-resolution microscope Elyra 7 was funded by the MRC through a capital equipment call. Cartoons in figs 1a, 5a,5c and 5h were created with BioRender.com FIB-SEM raw data *cam_hum-airway-14500* used in Fig.1B left, S1,S3,S4,S6,S7,S8 were prepared and acquired at the U. of Montreal EM imaging facility, region of Interest (ROI) sample provided by Vito Mennella (University of Cambridge), 3D segmentation by Aaran Vijayakumaran (University of Cambridge. FIB-SEM raw data *cam_hum-airway-14771-b* used in Fig 1B right, S1 were acquired at the EM imaging facility at the Brabraham Institute. ROI provided by Vito Mennella (University of Cambridge), sample provided by Chris O’Collaghan (UCL, GOSH), sample prepared by Nobu Morone (University of Cambridge), imaged by Kirsty MacLellan-Gibson (Brabraham), 3D segmentation by Aaran Vijayakumaran (University of Cambridge). FIB-SEM raw data *jrc_hum-airway-14953vc* used in Fig 2A,B, S5: Region of Interest (ROI) provided by Vito Mennella (University of Cambridge), sample provided by Nobu Morone and Sophie Breugesem, (University of Cambridge), prepared for imaging by Zhiyuan Lu (HHMI/Janelia/EMSR), with imaging by Wei Qiu (HHMI/Janelia/ FIB-SEM Technology) and Christopher K. E. Bleck (HHMI/Janelia/FIB-SEM Technology), post-processing by Eric Trautman, Michael Innerberger and Stephan Preibisch (HHMI/Janelia/SciCompSoft), block preparation imaging thru post processing facilitated by Aubrey Weigel (HHMI/Janelia/CellMap Project Team). 3D segmentation by Aaran Vijayakumaran (University of Cambridge).

The authors want to gratefully acknowledge Harald F. Hess and C. Shan Xu for their contributions to developing Enhanced Focus Ion Beam Scanning Electron Microscopy (eFIB-SEM) and driving the applications forward of this technology.

## Author contributions

**Conceptualisation:** VM, AV, CG

**Methodology:** AV, AA, CG, KN, AM, NM, KM-G, JL, EL, HHMI/Janelia, VM

**Investigation:** AV, CG, SYB, LRH, MD, KM-G, NT, EL, CO, HHMI/Janelia

**Visualisation:** AV, AA, CG, EL, VM

**Funding acquisition:** VM

**Writing – original draft:** VM, AV, CG

**Writing – review & editing:** VM, AV, CG, AM, KN, MD, EL, CO, SYB

**Competing interests:** Authors declare that they have no competing interests.

## Data and materials availability

jrc_hum-airway-14953vc

Resolution: 10×10×10 nm

Visualization link: https://openorganelle.janelia.org/datasets/jrc_hum-airway-14953vc

EM data DOI: 10.25378/janelia.25625346

Segmentation DOI: 10.25378/janelia.26035180

Presented in: Fig. 2A, B, S5

cam_hum-airway-14500 Resolution: 15x15x15 nm

Visualization link: https://openorganelle.janelia.org/datasets/cam_hum-airway-14500

EM data DOI: 10.25378/janelia.26035168

Segmentation DOI: 10.25378/janelia.26035174

Presented in: Fig. 1B left, S1, S3, S4, S6, S7, S8

cam_hum-airway-14771-b Resolution: 8x8x8 nm

Visualization link: https://openorganelle.janelia.org/datasets/cam_hum-airway-14771-b

EM data DOI: 10.25378/janelia.26039314

Segmentation DOI: 10.25378/janelia.26039317

Presented in: Fig. 1B right, S1

## Supplementary Materials

### Materials and Methods

#### Cell culture and ALI Differentiation

Human bronchial epithelial cells hTERT immortalized (BCi-NS1.1) (*55*) and human nasal epithelial cells (BMI-280) (*56*) were cultured in PneumaCult-Ex Basal Medium (STEMCELL Technologies) according to manufacturer instructions. After reaching 80% confluency, cells were detached by addition of Trypsin-EDTA (0.05 %) (Gibco) and seeded onto HTS Transwell Permeable Support plates with polyester membrane and 0.4 μm pore size, (Corning). Cells were grown until confluency for 3-4 days in PneumaCult-Ex Basal Medium (STEMCELL Technologies) before removal of apical media and differentiation in PneumaCult-ALI Medium (STEMCELL Technologies) for 28 days. All experiments except for Figs 1, 2a-c,f,g,i,l, 3, 4g, S2-S7, S8a,c-e,h, S11a,e were conducted with hTERT immortalized human bronchial epithelial cells.

#### Generation of CRISPR-Cas9 Knockout Cell Lines

*sgRNA design:* Guide RNAs targeting the human *CROCC* and *CCNO* genes were designed using the CRISPOR guide selection tool (*57*) using the GRCh38/hg38 genome and 20bp-NGG protospacer adjacent motif (PAM) sequence. Guide RNAs were selected to ensure a combined specificity score over 90, a predicted efficiency score over 55, and a predicted frameshift ratio over 75. Guides selected did not include any predicted off-targets with 2 bp mismatches, or any exonic 3 bp mismatches in the protospacer sequence. sgRNA sequences are: *CROCC* – GCUGCCGUUGGAAGCAUCCG; *CCNO* – GCUCUGGCCGUAGUCGCGGA; Scrambled – GCACUACCAGAGCUAACUCA

##### RNP complex generation

Synthetic modified sgRNA (2′-O-methyl analogs and 3′ phosphorothioate internucleotide linkages in first three nucleotide at both 5′ and 3′ ends) (Synthego) targeting *CROCC,* and *CCNO* loci were resuspended in nuclease-free 1X TE buffer alongside a scrambled control sgRNA. Ribonucleoprotein (RNP) complexes were made by 10 min RT incubation 1:1 with TrueCut Cas9 Protein v2 (Invitrogen) diluted in Buffer R (Invitrogen).

##### sgRNA in vitro cleavage assay

The target region for each sgRNA tested was amplified by PCR using Q5 High-Fidelity DNA Polymerase (New England Biolabs), purified (QIAquick Gel Extraction Kit, QIAGEN) and quantified (Nanodrop^TM^). sgRNAs (Synthetic modified sgRNA, Synthego) were resuspended in 1X TE buffer to 50 μM stock concentration. For each reaction, sgRNA (30 nM final) was set up 1:1 with Cas9 (TrueCut Cas9 Protein v2, Invitrogen) at room temperature (RT) with 1:10 v/v NEBuffer 3.1 before incubation at 37 °C for 10 min. Purified target DNA (3 nM final) was then added before further incubation at 37 °C for 1 hr. Reactions were terminated with 1 μl Proteinase K and 10 min RT incubation and loaded to 2 % agarose gel for fragment analysis by densitometry (Gel Doc EZ Imager, Image Lab Software, BioRad).

##### Electroporation & clonal selection

RNP complex was transfected into cell suspensions using NEON transfection system (Invitrogen) at a concentration of 6 pmol RNP complex per 100,000 cells. Transfected cells (including scrambled control) were cultured in PneumaCult-Ex Basal Medium (STEMCELL Technologies). Clonal populations of CROCC and CCNO KO BCi-NS1.1 cells were generated by limiting dilution. Isolated clones were expanded, cryopreserved, and validated.

##### gDNA Extraction and ICE Validation

Cells from each knockout pool, and scrambled control were pelleted (300 × g, 5 min) and gDNA extracted using GeneJET Genomic DNA Purification Kit (Thermo Scientific). Loci targeted by sgRNA were amplified from knockout pool and scrambled control gDNA by PCR using Platinum SuperFi II PCR Master Mix (Invitrogen) and products were purified (QIAquick PCR Purification Kit, QIAGEN) and analysed by Sanger sequencing. Knockout efficiency was determined by Inference of CRISPR Edits (ICE) analysis (Synthego). The result showed 96% indel, model fit of 97, and a knockout score of 96 for CROCC KO and 81% indel, model fit of 95, and 89% indel, model fit of 94, and knockout score of 85 for CCNO. CROCC KO clone 2_F10 observed genotype by sequencing was homozygous 1 bp insertion and showed 98% indel, model fit of 98, and a knockout score of 98 by ICE. CCNO KO clone 1_B07 observed genotype by sequencing was trans-heterozygous 2 bp deletion / 2 bp to 3 bp replacement (CG to GCC), and showed 96 % indel, model fit of 96, and knockout score of 96 by ICE.

#### Molecular Cloning

N-terminal eGFP-CROCC CDS (NM_014675.5) fusion was inserted into PiggyBac vector pPB-CAG-dmKO2-IN by amplification from pEGFP-CROCC/Rootletin construct (Addgene #41166) and restriction cloning. pEGFP-Rootletin was digested by *NotI* and *AseI* and pPB-CAG-dmKO2-IN was digested by *NotI* and *NdeI* (produces compatible cohesive ends with *AseI*) (37 °C, overnight). Enzymes were heat inactivated (65 °C, 20 min) and fragment cutting was confirmed by gel electrophoresis (1 % TAE agarose gel, 120 V, 1 hr). Fragments were excised and gel extracted (QIAquick Gel Extraction Kit, QIAGEN) followed by quantification (Nanodrop^TM^) and ligation 1:1 using T4 DNA ligase (16 °C, overnight). Each ligation was transformed into HB101 competent *E. coli* (Promega) followed by selection on ampicillin (100 μg/ml) LB plates. Resulting colonies picked and grown (37 °C, 180 rpm, overnight). Cultures prepped (Plasmid Mini Kit, QIAGEN) and quantified (Nanodrop^TM^). Validation of correct insertion by *EcoRI* digestion (37 °C, 1 hr) and gel electrophoresis (1 % TAE agarose gel, 120 V, 1 hr). PiggyBac vectors encoding either N-terminal or C-terminal eGFP-tagged CROCC2 (NM_001351305.2) under a CMV promotor with a Neomycin resistance cassette were designed and ordered from VectorBuilder. pPB-CAG-dmKO2-IN was a gift from Dr Feride Oeztuerk-Winder. pEGFP Rootletin (Nigg pFL2(CW499)) was a gift from Erich Nigg (Addgene plasmid # 41166; http://n2t.net/addgene:41166; RRID:Addgene 41166).

#### PiggyBac Transfection

hTERT immortalized human bronchial epithelial cells (BCi-NS1.1) at 80% confluency were detached by addition of Trypsin-EDTA (0.05 %) (Gibco). Transposon plasmids were electroporated into cell suspension alongside a hyperactive transposase vector pRP[Exp]-mCherry-CAG>hyPBase (VectorBuilder) using NEON transfection system (Invitrogen) using a ratio of 1:1 at 250 ng vector per 100,000 cells. The transposase mediates integration of the inverted terminal repeat sequences (ITRs) and cargo between these sites into TTAA chromosomal sites. Successfully transduced cells were cultured in PneumaCult-Ex Basal Medium (STEMCELL Technologies) and selected by G418 (20 μg/ml).

#### High-Resolution Respirometry

High-resolution respirometry of control and CROCC/Rootletin KO differentiated cells was carried out using the Oxygraph-2K system (Oroboros Instruments, Innsbruck, Austria). 1.5 million cells/ml were suspended in air-saturated mitochondrial respiratory medium (MiR05; 0.5 mM EGTA, 20 mM HEPES, 10 mM KH_2_PO4, 60 mM lactobionic acid, 3 mM MgCl_2_.6H20, 110 mM sucrose, 20 mM taurine, 1g L-1 bovine serum albumin, pH 7.1) at 37°C, and oxygen consumption recorded using DatLab v6.1. Basal level respiration was measured before titration with the mitochondrial uncoupler carbonyl cyanide p-trifluoro-methoxyphenyl hydrazone (FCCP) in 0.5 µM increments to reveal maximal electron transfer system capacity. Finally, the inhibitors antimycin A (2.5 µM) and sodium azide (100 mM) were added to reveal non-mitochondrial oxygen consumption, which was subtracted from other rates to give mitochondrial respiration.

#### Antibodies

See Table S1 for details of antibodies used in this study.

#### Immunofluorescence Labelling

Human airway epithelial cells were fixed in 4 % paraformaldehyde (PFA) for 30 minutes, washed three times and then blocked with 4 % BSA in PBS with 0.2 % Triton-X100 for 1 hr at room temperature (RT) after three washes. Alternatively, cells were fixed with −20°C histological grade anhydrous methanol (Fisher Chemical) for 30 min, washed three times and blocked with 4 % BSA and 0.05 % Tween-20 in PBS for 1 hr at RT. Primary antibodies were incubated at RT for 1 hr or overnight at 4 ℃. For secondary antibodies, samples prepared for 3D-SIM^2^ nanoscale mapping and rotational polarity analysis experiments used Cy™3 AffiniPure-VHH™ Fragment Alpaca Anti-Mouse IgG (H+L) (1:200) to label the reference marker (POC1B) and Alexa Fluor® 488 AffiniPure-VHH™ Fragment Alpaca Anti-Rabbit IgG (H+L) (1:200) to label the various centriolar and rootlet markers. VHH secondaries were incubated for 30 min at RT. For other immunostaining experiments, Alexa-dye conjugated secondary antibodies (1:400 – 1:500) were used and incubated for 1 hr at RT. All washing steps were performed with 1X PBS for PFA fixed samples and 0.05 % Tween-20 in PBS for methanol fixed samples for 7 minutes each. Cells on #1.5 high precision glass coverslip were mounted with ProLong Gold Antifade mounting medium (Invitrogen) and sealed with nail polish on glass slides. For STED experiments, Anti-mouse or anti-rabbit conjugated with Abberior STAR RED or STAR ORANGE dyes were used at 1:200 and incubated for 1 hr at RT. For 2D STED, the cells were mounted with Abberior Mount Solid Antifade.

For immunofluorescence of human airway epithelial cells on the side for cilia length quantification and assessment of mitochondrial positioning and distribution, differentiated cells were detached by addition of Trypsin-EDTA (0.05 %) (Gibco) and the cells were centrifuged at 500 x g for 5 min. The resulting pellet was resuspended in 4 % PFA for 20 min, washed three times with 1X PBS, and then blocked with 4 % BSA in PBS with 0.2 % Triton-X100 for 30 min. Alpha-tubulin FITC (1:400) and directly labelled TOM20 (1:400) were incubated at RT for 1 hr, and then washed three times with 1X PBS. After washing and antibody incubation steps, the cells were centrifuged at 500 x g for 5 min. After immunofluorescence staining, the resulting pellet was resuspended in 15 μL of ProLong Gold Antifade mounting medium (Invitrogen), placed onto a slide and covered with a #1.5 high precision glass coverslip.

#### Super Resolution Fluorescence Microscopy

##### 3D-Structured Illumination Microscopy (SIM)

Micrographs were acquired using a ZEISS Elyra 7 system (Carl Zeiss Microscopy) equipped with Plan-Apochromat 63x/1.40 Oil DIC f/ELYRA objective or Plan-Apochromat 40x/1.40 Oil DIC M27 (for fig. 5e) and Duolink 4.2 CL HS pco.edge sCMOS cameras. The fluorophores were excited with lasers wavelengths: 405 (500 mW) 488 (500 mW); 561 (500 mW), 647 (500 mW). The emission wavelengths were collected by passing through band-pass filter 495-550, long-pass 655 nm. For each individual z slice, thirteen raw images were acquired to collect phase information for Lattice SIM reconstruction.

For tile scan imaging (for fig. S14), apotome mode was used and micrographs were acquired using the Plan-Apochromat 25x/0.8 Imm Kor DIC M27 objective and Duolink 4.2 CL HS pco.edge sCMOS cameras. For each individual z slice, five raw images were acquired to collect phase information for Lattice SIM reconstruction. Raw data were reconstructed using the SIM or SIM^2^ module of ZEN Black 3.0 software version 16.0.18.306 with the auto sharpness filter function. Channel alignment was performed using a .bin calibration file generated from data collected with Tetraspec beads calibration slide (Carl Zeiss Microscopy).

##### 2D Stimulated Emission Depleted Microscopy (STED)

Micrographs were acquired with the STEDYCON (Abberior Instruments) equipped with a 60X NA1.4 Oil objective. Browser-based control software (Abberior Instruments) was used to generate the STED images. Images were acquired with a pinhole size of 0.70 aU and a pixel size of 15 nm. Excitation lasers for STAR ORANGE and STAR RED were run at 1-10 % and depleted with the STED laser at 50 % and 45 %, respectively. Signals were detected with a gate of 8 ns.

##### Airyscan Imaging

Micrographs were acquired using the Confocal ZEISS LSM 980 multiphoton inverted with AiryScan2 (Carl Zeiss Microscopy). Briefly, labelled samples were sequentially excited with laser lines at 639 nm, 561 nm, 488 nm, and 405 nm. Emission fluorescence was collected using a C Plan-Apochromat 63x/1.40 oil immersion objective (Carl Zeiss Microscopy) with an open pinhole and passed through an appropriate custom bandpass filter based on the expected emission profile of the sample to the arrayed detector for the Airyscan unit. Airyscan joint deconvolution was settings used was 10 iterations.

#### Lattice SIM^2^ Live-Cell Imaging

For live-cell imaging of eGFP-CROCC expressing multiciliated cells, the epithelia were separated from plastic transwells by cutting edges of the transwell membrane with a razor blade and placed face down on a µ-Dish 35 mm, high glass bottom imaging dish (Ibidi). Cells were covered with a 10 μL of FluoroBrite™ DMEM (ThermoFisher Scientific); a small weight was placed on top of the membrane to ensure it was flat during imaging. Live cell imaging was performed on ZEISS Elyra 7 system in incubation chamber at 37 °C with 5 % CO2 and imaged with 4.2 CL HS pco.edge sCMOS camera using 488 nm laser line, with Plan-Apochromat 63x 1.4 NA oil objective. For each z plane, 13 phases at 1 ms exposure time were acquired. Raw videos were reconstructed using the SIM^2^ module of the ZEN Black 3.0 software version 16.0.18.306.

#### FIB-SEM Sample Preparation, Acquisition, and 3D Segmentation

The *cam_hum-airway-14500* sample and FIB-SEM volume was collected and processed as previously described (*36*). Other differentiated human airway cells cultures grown on ALI filters (*cam_hum-airway-14771-b* and *jrc_hum-airway-14953vc*) were fixed with Karnovsky’s fixative in 0.1 M sodium cacodylate buffer (SCB, pH 7.4) with 0.05 % malachite green oxalate salt (M9015, Sigma-Aldrich Inc., MO USA) and processed largely as previously described (*58*,*59*). Briefly, specimens were post-fixed with the osmium-thiocarbohydrazide-osmium (OTO) method, with incubations in 1 % osmium tetroxide in SCB for 1.5 hr at RT, 2.5 % potassium ferrocyanide in SCB for 1.5 hr at RT, aqueous 1 % thiocarbohydrazide solution for 45 min at 40 °C, and then with a 2 % osmium tetroxide aqueous solution for 1.5 hr at RT, followed by en-bloc staining with aqueous 1 % uranyl acetate solution overnight at 4 °C. Samples were subsequently transferred in lead aspartate solution (pH 5.0) for 2 hr at 50 °C. After dehydration in ethanol/acetonitrile series, the sample was embedded in Epon/Araldite mixture. To prepare for sandwich-embedding, slide-glasses were treated with Sigmacote (Sigma SL2) to facilitate easy detachment of embedded cultures from the ALI filter post-polymerisation, and then dissected using ophthalmic microscalpels. The target regions were trimmed-down, mounted on aluminium pin stubs with conductive epoxy (CircuitWorks CW2400) and first imaged by SBF-SEM in FEI Quanta FEG 250 (Thermo Fischer Scientific, Einthoven Netherlands) equipped with ‘3View2XP’ system (Gatan Inc, Pleasanton, USA) as described previously (60) to check for sample quality.

For imaging of the *cam_hum-airway-14771-*b sample, specimens were placed into a Zeiss Crossbeam 550 scanning electron microscope running Zeiss Atlas (Zeiss, Oberkoch, Germany; version 5.3.5.3). A region of interest was located in SEM mode by imaging at 10 kV. The area was prepared for imaging by depositing tracking marks and milling a trench with the FIB beam to allow SEM imaging of the milled face. Electron micrographs were obtained with isotropic 10 nm resolution with a 9 µs dwell time. The SEM was operated at 2 kV accelerating voltage and 500 pA current. Both backscatter and secondary electron data were collected, to improve signal to noise ratio. Ion beam milling was performed at an accelerating voltage of 30 kV and current of 700 pA at a rate of 15.9 nm per minute. Image stacks were initially aligned using the Atlas 5 software, before the combined image channels were exported as individual .tiff files.

Images were acquired as a series of z-slices and were aligned using Etomo, the graphical user interface (GUI) to the IMOD package (*60*). 3D segmentation of cilia, ciliary rootlets, endosomal and autophagic organelles, cellular membranes, vesicles and lipid droplets from *cam_hum-airway-14500* and *cam_hum-airway-14771-b* and the mitochondria from *cam_hum-airway-14771-b* were performed manually with 3Dmod within the IMOD package (*60*,*61*). Manual segmentation was aided with interpolation at approximately every tenth slice to ensure continuity. The slicer dialog within 3Dmod was used for segmentation of the ciliary rootlets, to aid and guide accurate segmentation. 3D segmentation of the lipid droplets, nucleus, autophagic organelles, mitochondria Golgi bodies in *cam_hum-airway-14771-b* and *jrc_hum-airway-14953vc*, a deep-learning-driven approach was applied using the Empanada-Napari Plugin (*14*). To build image models for automated segmentation, a training set was built by manually segmenting patches (256 x 256 pixels each) containing the organelles of interest. To improve model accuracy, the fine-tuning module was used. After 3D inference on the entire volume, to display segmentation of the organelles within the cell of interest, we developed a Python script (available at GUI) that uses the mask of the target cell membrane generated with 3Dmod to extract organelles segmented within the specified cell/mask. After extraction, the segmentation masks were proofread to ensure accurate annotation. The final segmentation masks were binned by three or four depending on the size of the original segmentation mask, converted to binary .tiff file stacks and visualised in 3Dmod using the isosurface rendering dialog function.

#### FIBSEM Datasets

Below is a brief description for each dataset used in this work. The volumes are stored on the cloud using Amazon Web Services (AWS) S3 storage platform and viewable via OpenOrganelle: openorganelle.janelia.org.

##### cam_hum-airway-14500

Contributions: Samples were prepared and acquired at the U. of Montreal EM imaging facility, region of Interest (ROI) sample provided by Vito Mennella (University of Cambridge), 3D segmentation by Aaran Vijayakumaran (University of Cambridge).

Voxel size: 10 x nm 10 nm x 10 nm

Visualization link: http://janelia-openorganelle-web-dev.s3-website-us-east-1.amazonaws.com/datasets/cam_hum-airway-14500

EM data DOI: 10.25378/janelia.26035168

Presented in: Fig. 1B left, S1, S2, S3, S4, S6, S7, S8

##### cam_hum-airway-14771-b

Contributions: Samples were prepared by Nobu Morone (University of Cambridge) and acquired by Kirsty MacLellan-Gibson at the EM imaging facility at the Babraham Institute. ROI provided by Vito Mennella (University of Cambridge), sample provided by Chris O’Collaghan (UCL, GOSH), 3D segmentation by Aaran Vijayakumaran (University of Cambridge).

Voxel size: 10 x nm 10 nm x 10 nm

Visualization link: http://janelia-openorganelle-web-dev.s3-website-us-east-1.amazonaws.com/datasets/cam_hum-airway-14771-b

EM data DOI: 10.25378/janelia.26039314

Presented in: Fig. 1B right, S1, S2

##### jrc_hum-airway-14953vc

Contributions: Samples were prepared for imaging by Zhiyuan Lu (HHMI/Janelia/EMSR), with imaging by Wei Qiu (HHMI/Janelia/ FIB-SEM Technology) and Christopher K. E. Bleck (HHMI/Janelia/FIB-SEM Technology), post-processing by Eric Trautman, Michael Innerberger and Stephan Preibisch (HHMI/Janelia/SciCompSoft), block preparation imaging thru post processing facilitated by Aubrey Weigel (HHMI/Janelia/CellMap Project Team), samples provided by provided by Nobu Morone and Sophie Breusegem, (University of Cambridge), Region of Interest (ROI) provided by Vito Mennella (University of Cambridge), 3D segmentation by Aaran Vijayakumaran (University of Cambridge).

Voxel size: 8 x nm 8 nm x 8 nm

Visualization link: http://janelia-openorganelle-web-dev.s3-website-us-east-1.amazonaws.com/datasets/jrc_hum-airway-14953vc

EM data DOI: 10.25378/janelia.25625346

Presented in: Fig. 2A, B, S5

#### High-pressure Freezing and Freeze-substitution TEM

Human airway epithelium cells were differentiated on transwells covered with Quantifoil UltraAuFoil grids (R1.2/1.3 gold 300-meshes, Quantifoil Micro Tools GmbH, Germany) that were glow-discharged (GloQube, Quorum Tecknologies, East Sussex UK) and treated with collagen solution. After 28 days of differentiation, cells on grids were used for high-pressure freezing and freeze substitution by using Leica EM-HPM100 and EM-AFS2/FSP (Leica Microsystems, Vienna Austria). Briefly, the specimens were kept in 0.05 % tannic acid in anhydrous acetone at −90 °C for at least 20 hr, washed with pure acetone at −90 °C, and substituted in 1 % osmium tetroxide, 0.1 % uranyl acetate, and 1 % pure water in acetone, with temperature controlled under −90 °C for 48 hr, at −60 °C for 8 hr, at -30 °C for 8 hr with interval of 30 °C/hr. After several washes with pure acetone at 4 °C, the airway cells were infiltrated for a few days and embedded in Epon and Araldite mixture (TAAB Laboratories Equipment Ltd., Reading UK). After polymerization at 65 °C for a few days and removal of the gold grid-mesh, ultrathin-sections (∼60 nm) were obtained with a Ultramicrotome (Leica EM-UC7/Artos-3D), mounted on formvar-carbon films/EM grids, stained with lead citrate and uranyl acetate, and then observed with a 200 kV transmission electron microscope (FEI TalosF200C, Oregon USA) with Ceta-16M CMOS-based camera (4kx4k pixels under 16-bit dynamic range), and JEM1400flash (JEOL, Tokyo Japan) with TemCam-XF4016 (TVIPS GmbH, Germany).

#### Bioinformatics

For single-cell RNA-seq (sc-RNAseq) analysis, data was downloaded from the Integrated Human Lung Cell Atlas (HLCA) core, which includes data of healthy lung tissue from 107 individuals (*62*) via cellxgene (https://cellxgene.cziscience.com/collections/6f6d381a-7701-4781-935c-db10d30de293). Data analysis was performed in R (version 4.3.0) using Seurat (version 5.0.1). Epithelial cell types of interest – ‘Multiciliated (nasal)’, ‘Basal resting’, ‘Deuterosomal’ and ‘Suprabasal’ were subsetted from the downloaded dataset. Next, a pseudobulking approach to aggregate the single-cell expression data for each cell type using the ‘AggregrateExpression()’ function, thereby creating bulk-like expression profiles. This helped to reduce the intrinsic noise and variability associated with individual cell expression measurements. Log normalisation of the expression data was then performed post-pseudobulking with ‘normLibSizes()’ function, to standardize gene expression levels across cells. Heatmaps were generated using the pheatmap package, with a focus on raw scaling to standardize expression data for each gene across cell types.

#### 3D segmentation Image Analysis

3Dmod Object Info function was used to quantify the volume and surface area of segmented ciliary rootlets. Fiji free-hand drawing tool was used to measure rootlet length. Morphometric analysis of the mitochondria was performed by first extracting individual mitochondria from the reconstructed volumes— selection bias was minimized by using a random number generator to determine the samples for extraction. The resulting image stacks, saved as .tif files, were then imported into ImageJ, transformed into a binary mask and processed with the ‘fill holes’ tool to facilitate subsequent analyses. The 3D Suite Plugin within Fiji was then applied to assess quantitative morphological parameters, including sphericity and axial elongation (*63*). For the analysis of ciliary rootlet-organelle interactions, proximity within 1 pixel (equivalent to 10 nm) between the rootlet and organelles was manually annotated. This analysis was facilitated by slicer module, complemented by the ‘Measure’ tool available within the 3Dmod.

#### Super-Resolution Nanoscale Mapping and Rotational Polarity Analysis

For 3D-SIM^2^ nanoscale mapping experiments, centriolar marker POC1B was used as reference marker in the 555 channel, relative to other markers in the 488 channel to ensure highest resolution. To ensure maximum distance accuracy, 3D-SIM^2^ micrographs where the fluorescence maxima of POC1B and the target marker appeared on the same z-plane were included for analysis. Distances between the fluorescence peak maxima of POC1B and the respective reference protein were determined using the built-in calliper function of the Zeiss Zen Blue imaging software.

To measure CROCC N-N and C-C termini distances, individual ciliary rootlets were extracted from 2D STED images, followed by analysis of fluorescence intensity profiles using Fiji’s ‘Plot Profile’ feature and custom Python script (available at GUI) employing the NumPy (Python 3.9.15)) and scipy.signal.find_peaks (Python 3.9.15) libraries to identify peaks indicative of termini positions within the intensity profiles. Consecutive peaks’ X-coordinates were computed to determine inter-termini distances and subsequently averaged to determine the periodicity.

Rotational polarity analysis was assessed using a modified version of a previously published MATLAB script (https://github.com/liuzhorizon/PCDDiagnosis_Quantitative_SuperResolution) (*32*). Briefly, cells were cultured on transwell filters under air-liquid interface conditions for 28 days until they were fully differentiated. Following fixation in −20 °C anhydrous methanol, cells were stained with antibodies against POC1B and centriolin to mark basal bodies and basal feet, respectively. 3D-SIM^2^ images were acquired and analysed by thresholding and segmentation to identify basal body and basal foot pairs, with a nearest neighbour cut-off distance 400 nm.

#### Cilia Length Quantification

To quantify cilia length within control and CROCC/Rootletin KO differentiated epithelial cells, cilia lengths were measured manually using the freehand line tool in the image processing software Fiji ImageJ. Only cilia with visible starting and end point from multiciliated cells on their side were measured.

#### Cilia Beat Frequency Analysis

For cilia beat frequency (CBF) measurements, transwell membranes from differentiated airway epithelia of control and CROCC KO cells were excised with a razor blade and placed face down on a µ-Dish 35 mm, high glass bottom imaging dish (Ibidi). The cells were covered with 10 μL of PneumaCult-Ex Basal Medium (STEMCELL Technologies) and a small weight on top of the membrane to ensure proximity to the objective during imaging. Live imaging was performed on the ZEISS Elyra 7 system equipped with Plan-Apochromat 63x/1.40 Oil DIC f/ELYRA objective or Plan-Apochromat 40x/1.40 Oil DIC M27 using transmitted light illumination. The sample was placed in an incubation chamber at 37 °C with 5 % CO2 and imaged for a total of 2000 frames at 1 ms exposure time. Videos of airway cells top down and on side views were then imported into FIJI for CBF analysis using FreQ plugin (*64*). The average CBF of the regions of interests was measured from 3 independent biological replicates

#### Structural Modelling Using AlphaFold

The structures of proteins and protein complexes shown in figures S12 and S15 were modelled using a local installation of AlphaFold multimer v2.3.1 [11, 12] as well as the web server version of AlphaFold v3 recently release (*65*). Sequences of *Homo sapiens* CROCC, CROCC2, CEP250 and LRRC45 were used as inputs for the structure predictions. Predicted Alignment Error (PAE) plots were prepared using the PAE viewer [13]. PyMOL v. 2.5 (Schrodinger LLC, https://pymol.org) was used for preparing figures of protein structures.

#### Statistical Analysis

Data were analysed in Microsoft Excel and Prism software. Statistical tests, sample sizes and number of replicates are specified in figure legends. Differences were regarded as significant if p < 0.05.

**Table S1:** Primary Antibody and Dyes List. . * = 1:1000 was used for S8, whilst 1:500 was used for other figures.

|  |  |  |  |  |
| --- | --- | --- | --- | --- |
| CEP44 | ProteinTech | 24457-1-AP | Methanol | 1:50 |
| CEP135 | Thermo Fisher Scientific | PA5-36440 | Methanol | 1:50 |
| CEP250 C-Termini | Thermo Fisher Scientific | PA5-66730 | Methanol | 1:200 |
| CEP250 N-Termini | ProteinTech | 14498-1-AP | Methanol | 1:100 |
| CROCC C-Termini | Santa Cruz<br>Biotechnology | sc-74056 | Methanol | 1:100 |
| CROCC N-Termini | Atlas Antibodies | HPA021191 | Methanol | 1:100 |
| CROCC2 | Thermo Fisher Scientific | PA5-61802 | Methanol | 1:50 |
| POC1B (WDR51B) | Thermo Fisher Scientific | H00282809-B01P | Methanol | 1:100 |
| LRRC45 | Thermo Fisher Scientific | PA5-54777 | Methanol | 1:100 |
| POC1B | Thermo Fisher Scientific | PA5-24495 | Methanol | 1:100 |
| TOM20 | Abcam | ab221292 | PFA | 1:500,<br>1:1000* |
| Alexa Fluor™ Plus<br>405 Phalloidin | Thermo Fisher Scientific | A30104 | PFA | 1:300 |
| Centriolin (C-9) | Santa Cruz<br>Biotechnology | sc-365521 | Methanol | 1:100 |
| Acetylated-Tubulin | Merck | T7451 | PFA,<br>Methanol | 1:600 |
| PCM1 | Santa Cruz<br>Biotechnology | sc-398365 | Methanol | 1:100 |

**Table S2:** Nanoscale Mapping Results. * = distance from reference marker POC1B

| **Protein** | **Mean distance\***<br>**(nm)** | **Mean distance**<br>**STD** | **Number of**<br>**centrioles** | **Number of cells** |
| --- | --- | --- | --- | --- |
| CEP44 | 186.1 | 20.8 | 97 | 10 |
| CEP135 | 284.0 | 33.9 | 112 | 10 |
| CEP250 CTD | 316.0 | 20.2 | 129 | 14 |
| CEP250 NTD | 458.5 | 34.5 | 113 | 12 |
| CROCC CTD | 510.9 | 67.6 | 115 | 12 |
| CROCC NTD | 598.7 | 73.5 | 108 | 11 |

**Table S3:** Rotational Polarity Analysis Result.

| **Condition** | **Number of cells analysed** | **Number of independent samples** | **Mean vector length** | **Median vector length** | **Vector length STD** |
| --- | --- | --- | --- | --- | --- |
| Control | 372 | 3 | 0.4744 | 0.4809 | 0.1603 |
| CROCC KO | 375 | 3 | 0.3634 | 0.3404 | 0.1742 |

**Table S4:** High Resolution Respirometry Analysis Result. ETC – Electron Transport Chain; * = units are pmol O_2_/s/million cells

| **Condition** | **Mean basal level respiration\*** | **Mean maximum ETC\*** | **Basal level respiration STD** | **Maximum ETC STD** | **Number of independent samples** |
| --- | --- | --- | --- | --- | --- |
| Control | 18.4846 | 65.3456 | 2.6903 | 1.8069 | 3 |
| CROCC KO | 19.2892 | 55.5457 | 2.0823 | 1.7058 | 3 |

**Fig S1.**
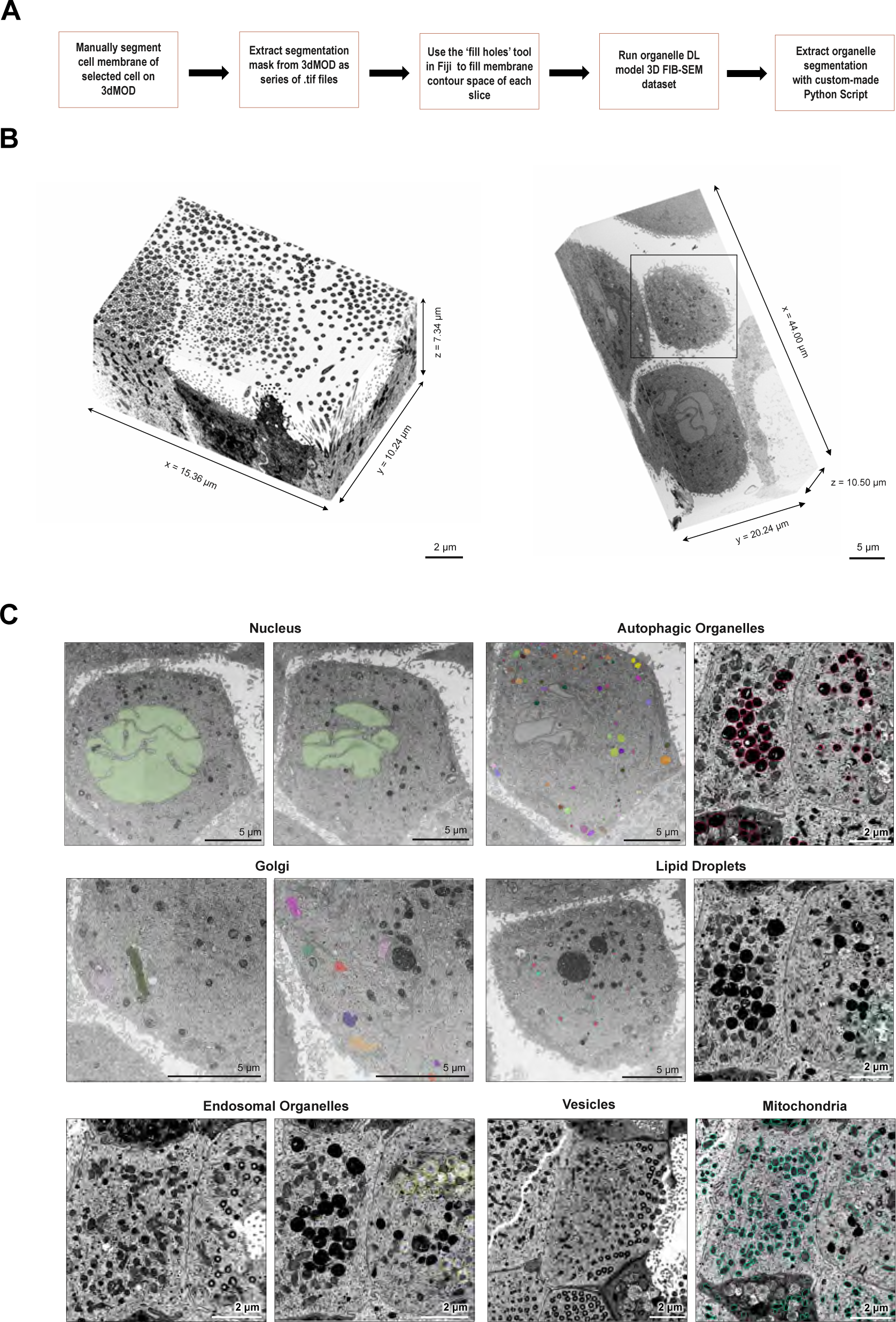
FIB-SEM 3D Volume Raw Data, Analysis and Organelle Segmentation. **(A)** Detailed workflow describing manual and deep-learning segmentation of FIB-SEM data with cell extraction to allow fast 3D reconstruction of airway epithelial cells that are amenable to single cell quantitative analysis. **(B)** Full raw volumetric data of human airway epithelium cells. The volume on the left contains airway multiciliated cells and a deuterosomal cell. The volume on the right contains the airway basal stem cell. **(C)** 2D FIB-SEM snapshots highlighting examples of segmented organelles from different volumes regions.

**Fig. S2.**
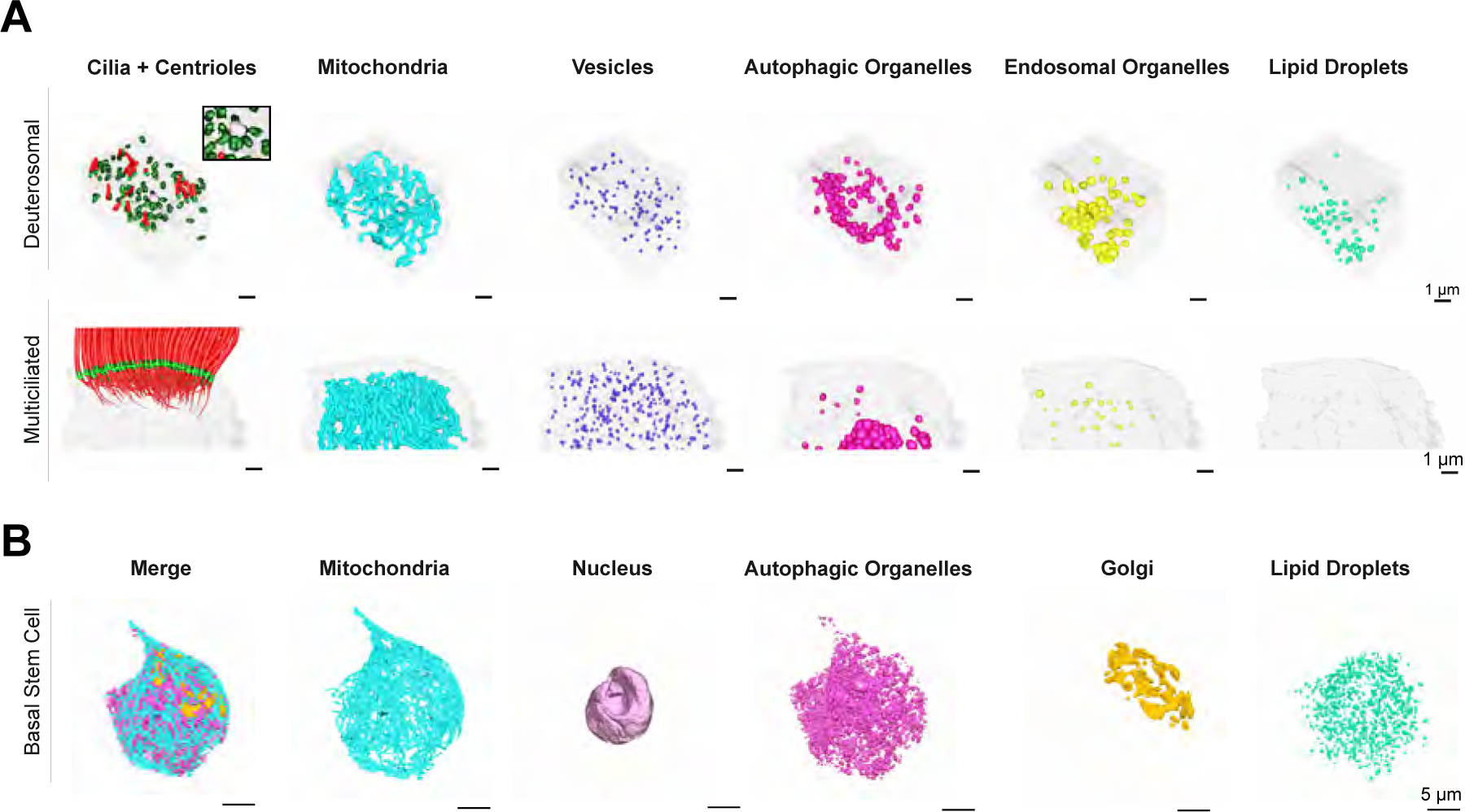
3D segmentation of individual organelles from airway epithelial cells. 3D segmentation of individual organelles in deuterosomal and MCC cells (A) or airway basal stem cell (B)

**Fig. S3.**
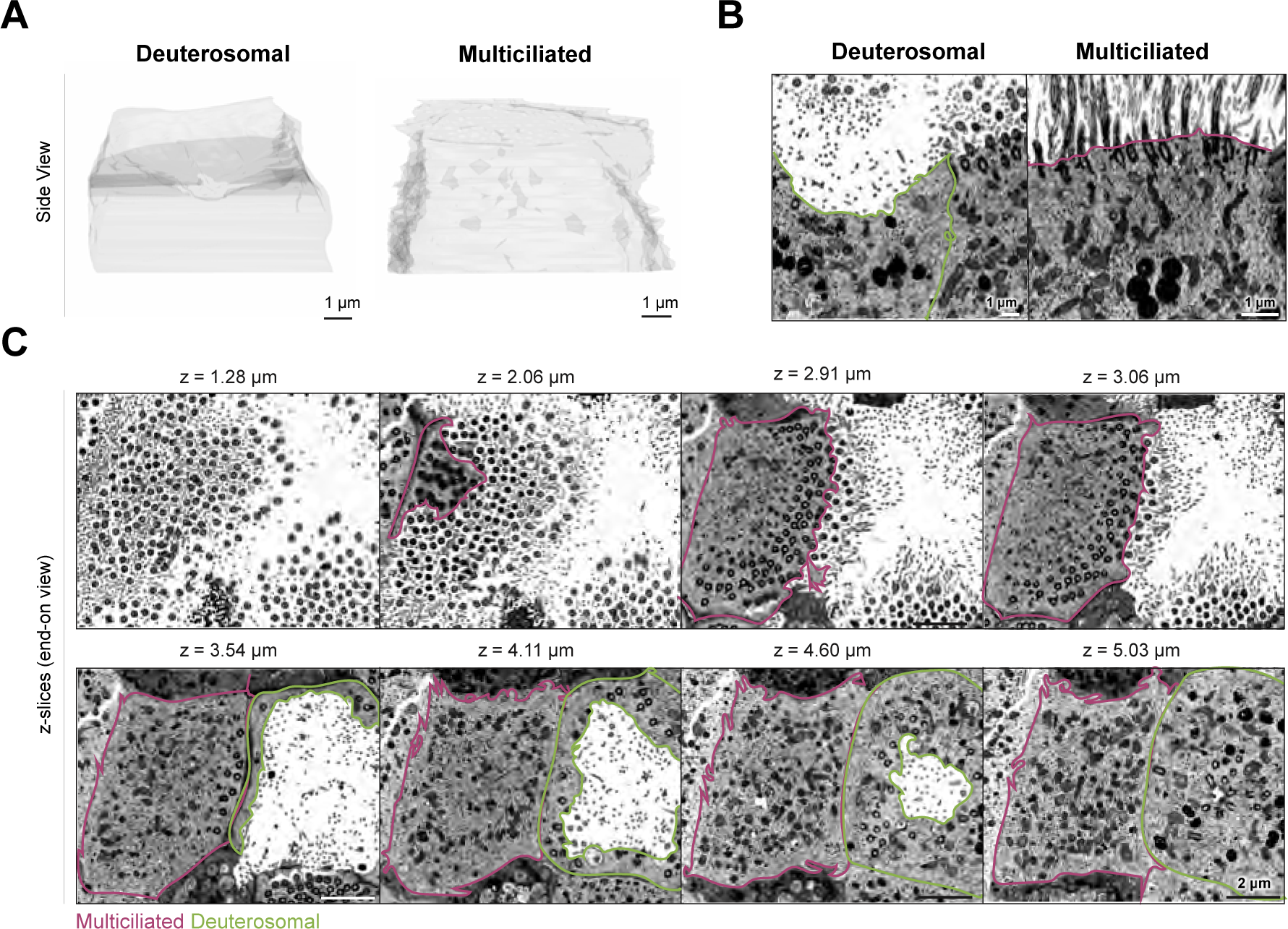
Plasma membrane organization of airway epithelial cells during multiciliogenesis. **(A)** 3D segmentation of plasma membrane on side view, highlighting the deuterosomal cell’s concave versus MCC’s convex surface. **(B)** 2D FIB-SEM slices on side view show differences in architecture of plasma membrane in the apical region of multiciliated (outlined in magenta) and deuterosomal (outlined in green). **(C)** Series of 2D FIB-SEM slices in end-on view highlighting differences in membrane topology between multiciliated (outlined in magenta) and deuterosomal cell (outlined in green).

**Fig. S4.**
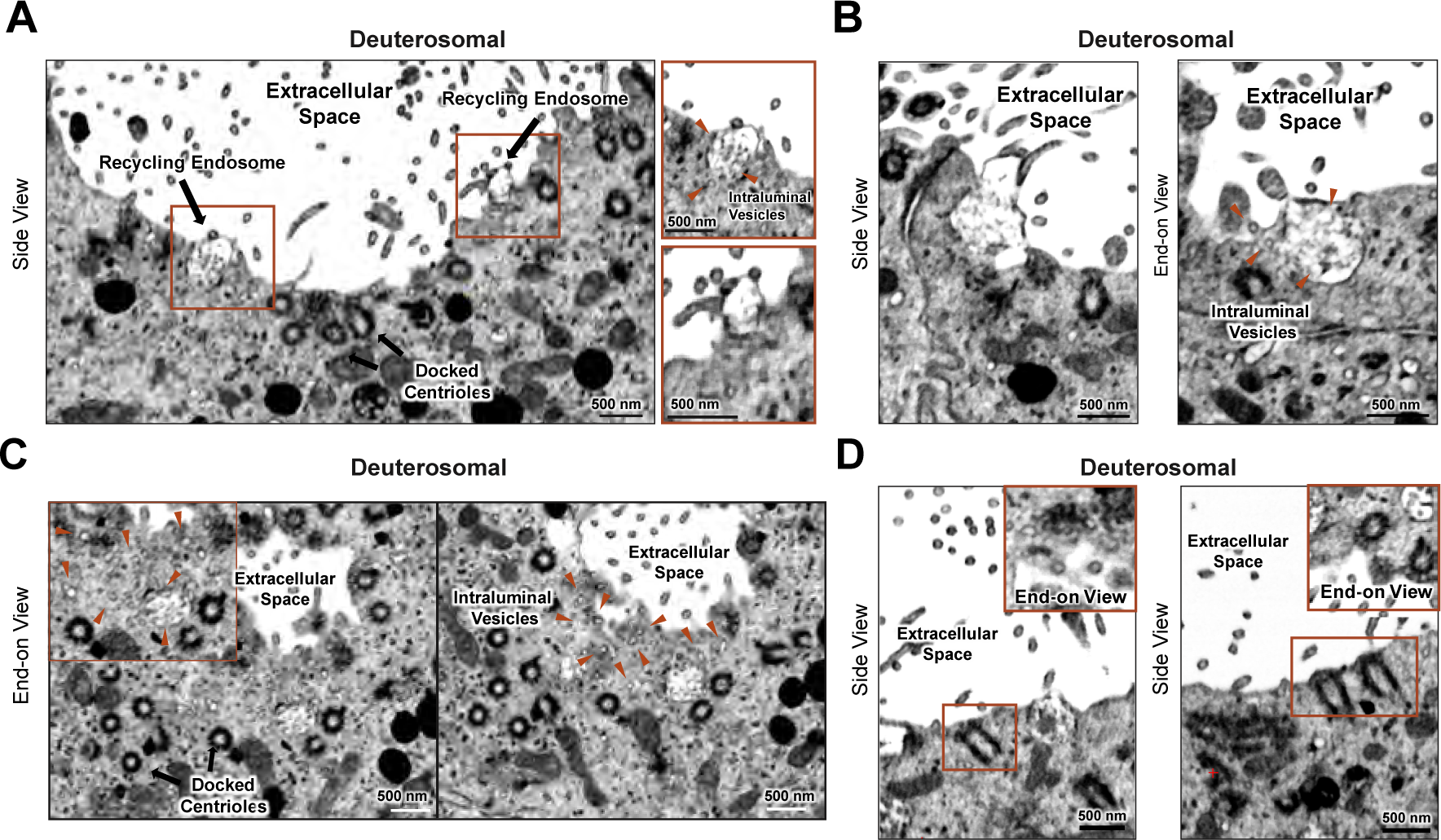
Deuterosomal cells present abundant fusion of recycling endosomes at the apical plasma membrane. **(A)** 2D FIB-SEM slices showing side views of docked centrioles to the apical membrane of the deuterosomal cell. Red insets highlight the fusion of apical recycling endosomes to the plasma membrane. Red arrowheads point to intraluminal vesicles within recycling endosome. **(B & C)** Examples of FIB-SEM slices showing apical recycling endosomes fusing with the plasma membrane in deuterosomal cells (red arrowheads). **(D)** FIB-SEM slices showing docked basal bodies lacking distinct ciliary vesicles.

**Fig. S5.**
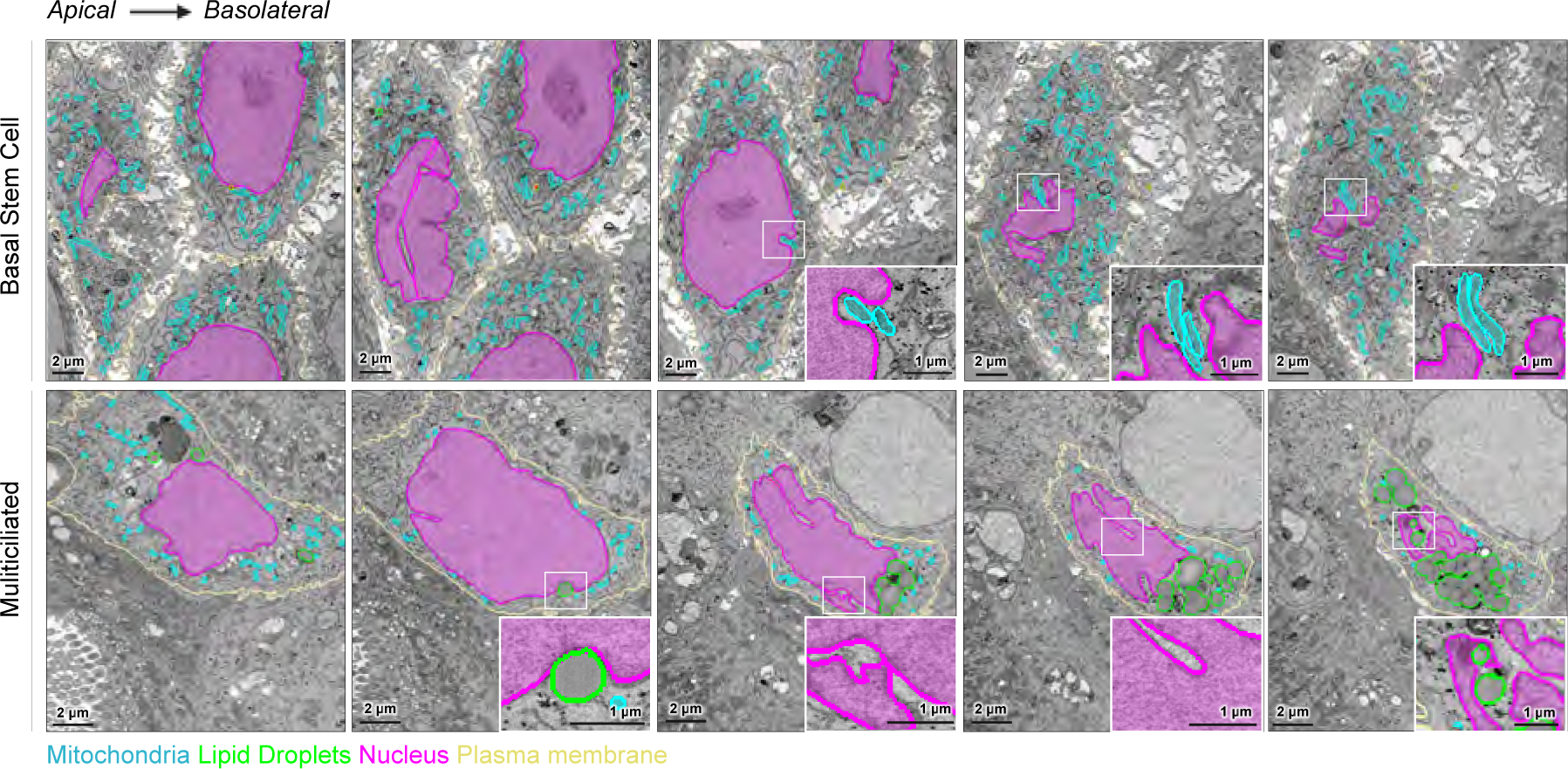
Perinuclear organelle composition changes during MCCs differentiation. **(A)** Series of 2D FIB-SEM slices in end-on view from apical to basolateral end of the pseudostratified epithelium (left to right) highlighting differences in organelle composition within nuclear invagination and around the nuclear envelope between airway basal stem cells and MCCs.

**Fig. S6.**
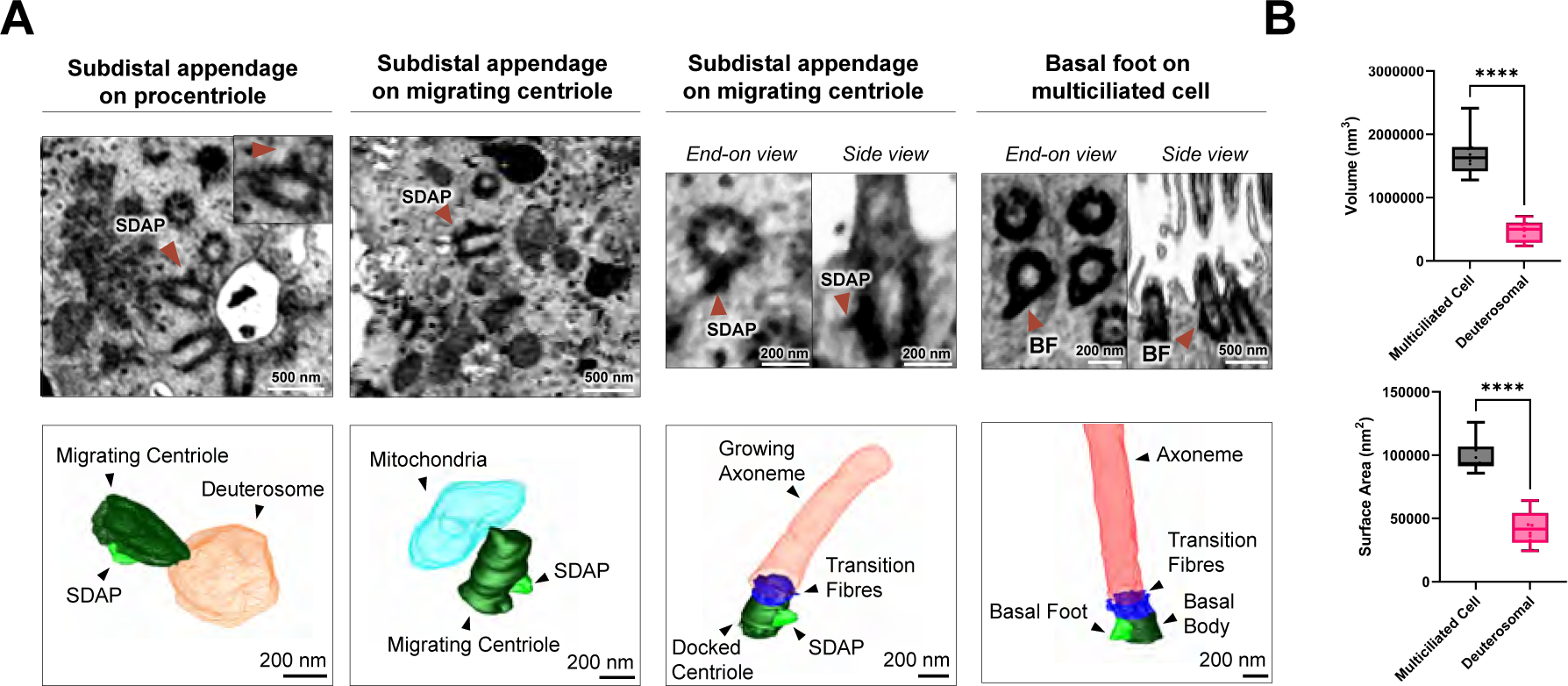
Structural transformation of subdistal appendages into basal feet during multiciliogenesis. **(A)** 2D FIB-SEM slices and corresponding 3D volume segmentation at different stages of MCC differentiation. From left-to-right, the first panel shows procentriole with a single subdistal appendage (SDAP) emanating from the deuterosome; the second panel shows a single SDAP on a migrating centriole; the third panel shows a SDAP on a docked centriole and an axoneme mid-growth; the fourth panel shows a mature basal foot (BF) on a docked centriole within a differentiated multiciliated cell (Deut, Deuterosome; SDAP, Subdistal Appendage; BF, Basal Foot). **(B)** Comparison of the volume and surface area of BF from differentiated MCC (mean volume = 1667000 nm^3^; mean surface area = 98986 nm^2^) or SDAP from deuterosomal cells (mean volume = 469551 nm^3^; mean surface area = 42125 nm^2^). **** p < 0.0001. (n = 10). Data were analysed with unpaired t-test.

**Fig. S7.**
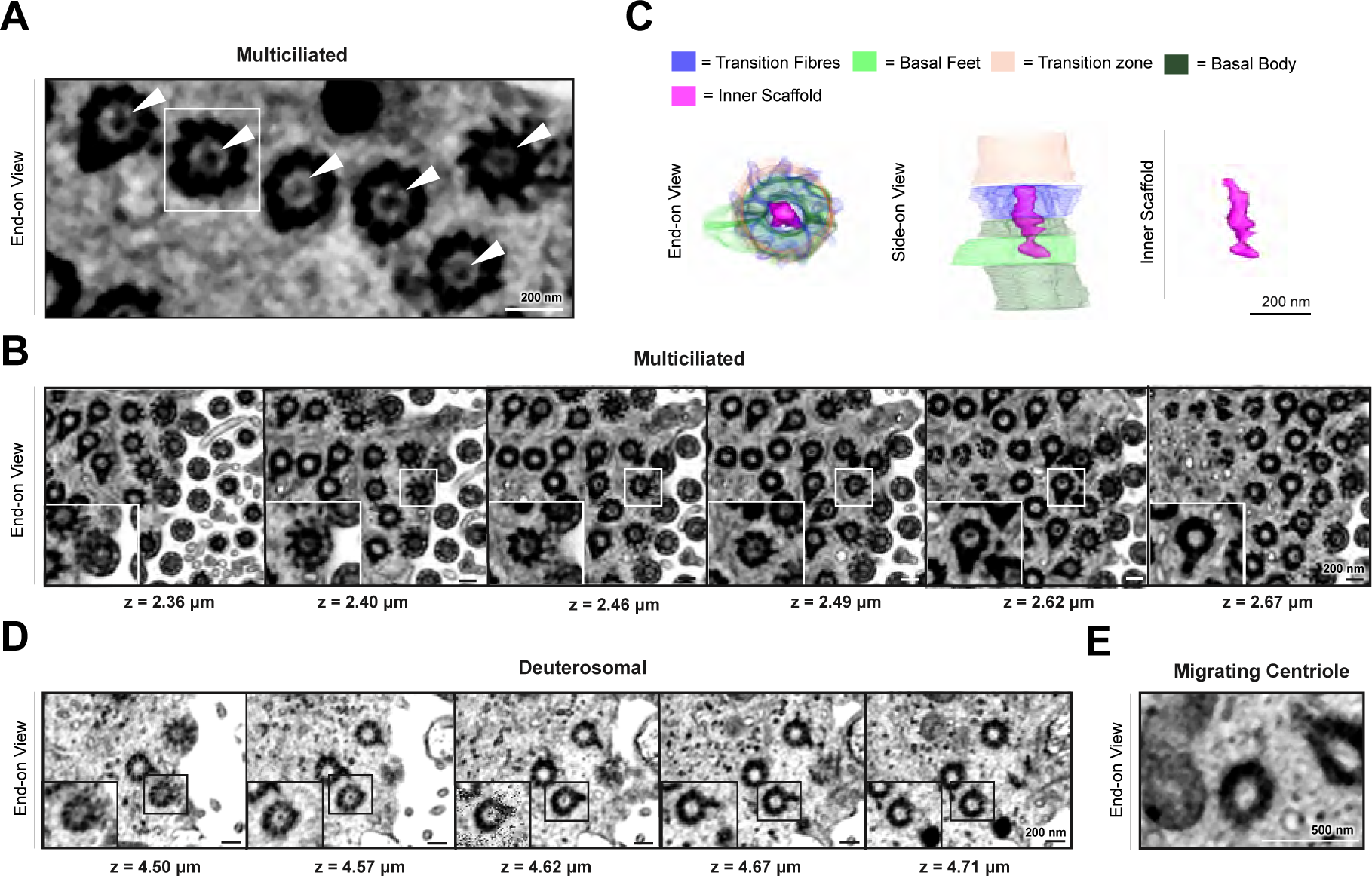
Basal bodies show internal scaffold in MCCs stage. **(A)** 2D FIB-SEM images of cross sections of basal bodies in a differentiated human airway MCC. Arrowheads indicate the density located in the distal region in the lumen of the basal body. **(B)** 2D FIB-SEM slices sequence highlights the length of the distal structure (from 2.36 μm to 2.87 μm). **(C)** 3D segmentation depicting the basal body (dark green), transition fibres (blue), basal feet (light green), transition zone (orange), and the filamentous distal structure (magenta). **(D)** 2D FIB-SEM slice showing a migrating centriole within a deuterosomal cell. Note the absence of the distal structure. **(E)** 2D FIB-SEM slices sequence showing a docked centriole in the deuterosomal cell. High magnification of the inset region highlights the absence of the dense distal structure observed in differentiated MCCs, except for a small density at z = 4.57 μm and 4.62 μm.

**Fig. S8.**
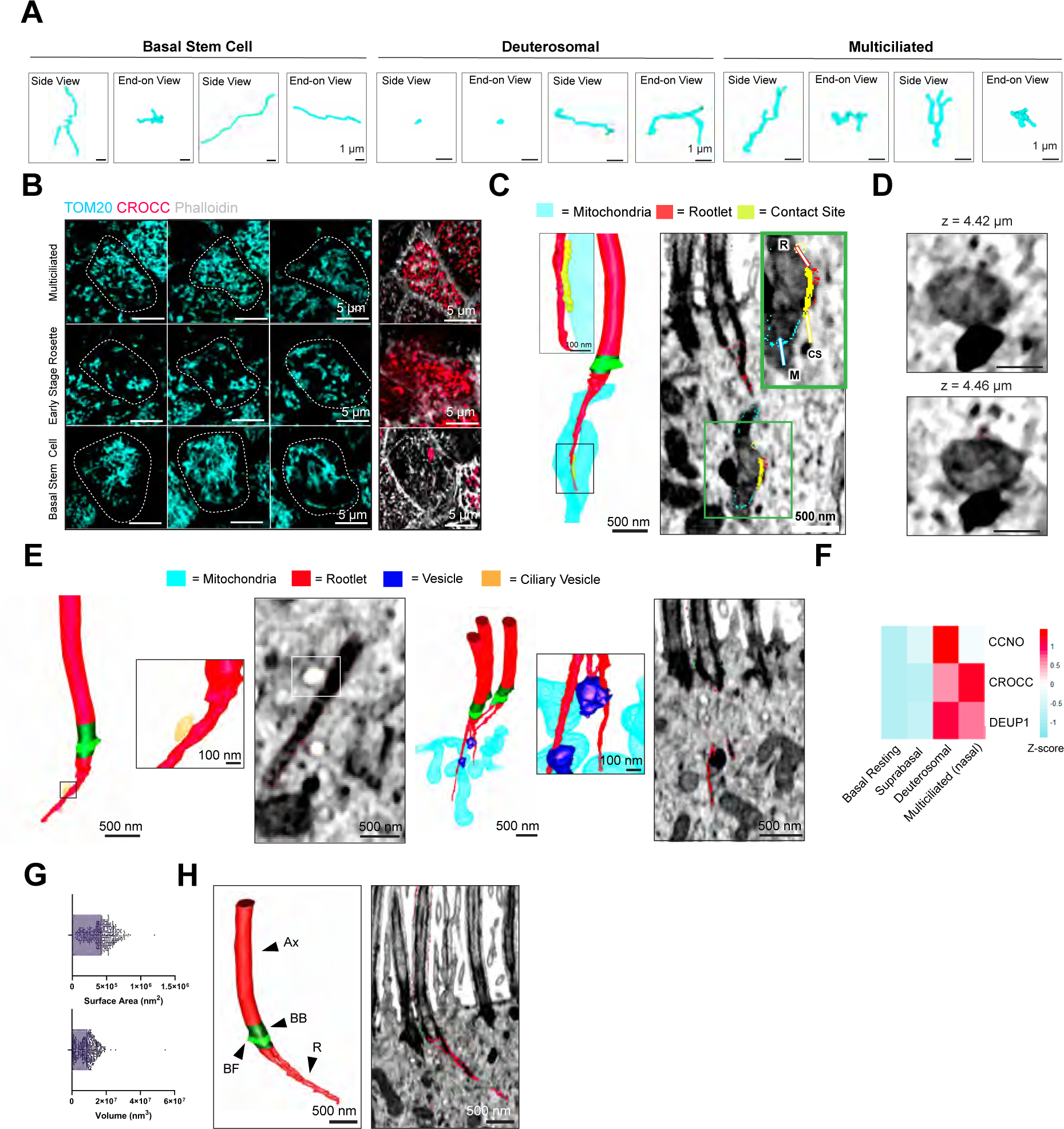
Mitochondrial organization and organelle-cytoskeleton interactions during MCCs differentiation. **(A)** Three-dimensional reconstructions of representative individual mitochondrion from the deuterosomal cell, differentiated MCC and airway basal stem cell state. **(B)** Airyscan fluorescence microscopy micrographs of different z sections showing organisation of mitochondria labelled with antibody anti-TOM20 (cyan) at different stages of differentiation in human airway epithelial cells. Right panel shows maximum intensity projection of phalloidin F-Actin labelling (white) used to detect cell boundaries and CROCC (red) to detect differentiating cells. **(C)** 3D volume segmentation of a rootlet (red) in contact with mitochondria (blue). Inset show high-magnification of the continuous interaction between the rootlet and mitochondria (yellow). The right panel shows corresponding 2D EM slices overlaid with segmentation contours. **(D)** Top panel: Corresponding series of EM slices (xy view) from (D) highlighting the rootlet-mitochondria interaction. **(E)** 3D volume segmentation of rootlet-organelle interactions: rootlet (red) and ciliary vesicle (gold); rootlet (red) and vesicle (dark blue). **(F)** Expression differences (log2-fold changes) of DEUP1, CCNO and CROCC in different cell types across human airway multiciliated differentiation from the Integrated Human Lung Cell Atlas (HLCA) study after pseudo-bulking. **(G)** Distributions of the rootlet surface area and volume, n = 412. **(H)** 3D segmentation of a branching rootlet and corresponding FIB-SEM slice overlaid with segmentation contours shown in movie 3. Notations: Ax (Axoneme), BB (Basal Body), BF (Basal Feet), R (Rootlet).

**Fig. S9.**
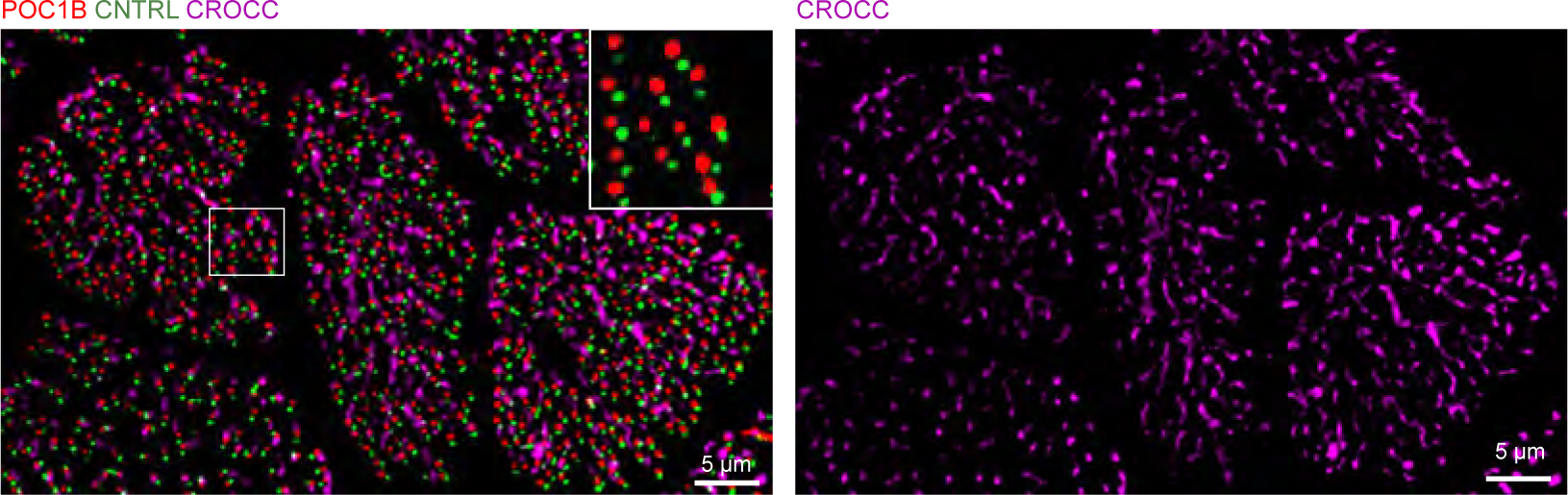
Rootlets cellular architecture by super-resolution microscopy. 3D-SIM volume maximum intensity projection of differentiated human MCCs (end-on-view). The left panel shows a merge projection of MCCs labelled with Centriolin, POC1B and CROCC showing the basal feet (green), basal body (red) and rootlets (merge) respectively. Inset shows high magnification highlighting aligned basal body/basal feet pairs. The right panel shows just the rootlets (CROCC) channel, delineating an asymmetrical dome structure skewed towards the posterior end of the cell relative to the effective stroke.

**Fig. S10.**
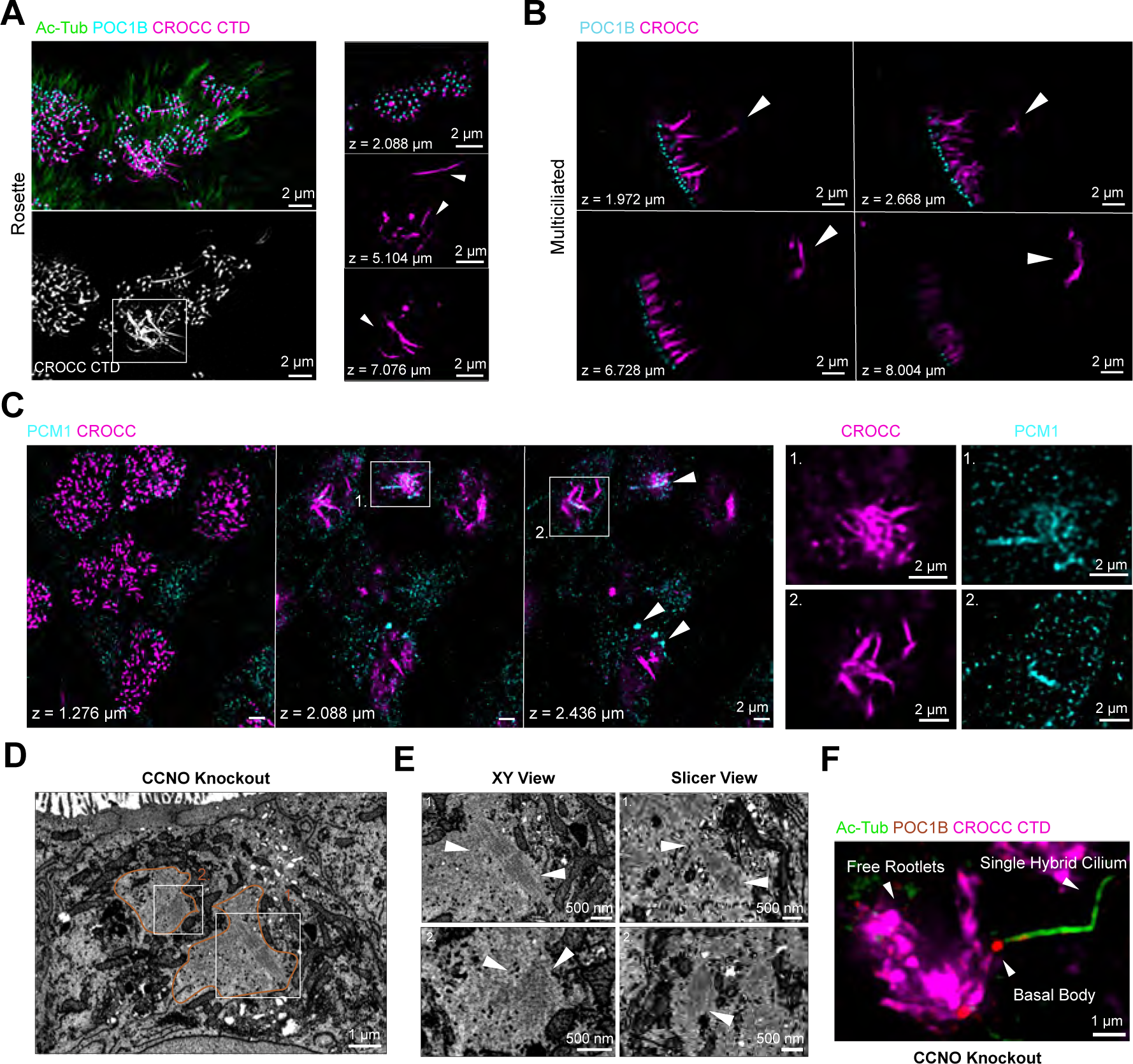
Multiciliated cells present free unattached rootlets. **(A)** Maximum intensity projection of 3D-SIM^2^ volume of partially differentiating human MCC at the rosette stage labelled with anti-POC1B (red), Ac-Tub (green), and CROCC CTD (magenta). Inset shows 3D-SIM^2^ single z-slices of the boxed region labelled with anti-POC1B (red) and CROCC CTD (magenta) antibodies. White arrowheads point at free cytoplasmic rootlets. **(B)** Maximum intensity projection of 3D-SIM^2^ volume of human MCCs, labelled with anti POC1B (cyan), Ac-Tub (green), and CROCC CTD (magenta) antibodies. White arrowheads point at free cytoplasmic rootlets. 3D-SIM^2^ z-slice series of a differentiated MCC labelled with anti POC1B (cyan), and CROCC CTD (magenta). White arrowheads point at free cytoplasmic rootlets. **(C)** Airyscan micrograph single z-slices of MCCs labelled with anti-CROCC (magenta) and anti-PCM1 (cyan). White inset boxes are magnifications of free rootlets at z = 2.088 μm (magnification 1) and z = 2.490 μm (magnification 2), showing colocalization of free rootlets and PCM1 at the basolateral end of the cell. **(D)** Single EM slice of an airway epithelial cell from CCNO KO cell line (side view) showing the presence of free rootlets within regions devoid of organelles (outlined in orange). **(E)** High-magnification view of boxed area in (D). slicer view of the volume was obtained using the slicer module within 3Dmod. **(F)** Maximum intensity projection of 3D-SIM volume from human airway cells CCNO KO labelled with anti-POC1B (red), Ac-Tub (green), and CROCC CTD (magenta) antibodies POC1B.

**Fig. S11.**
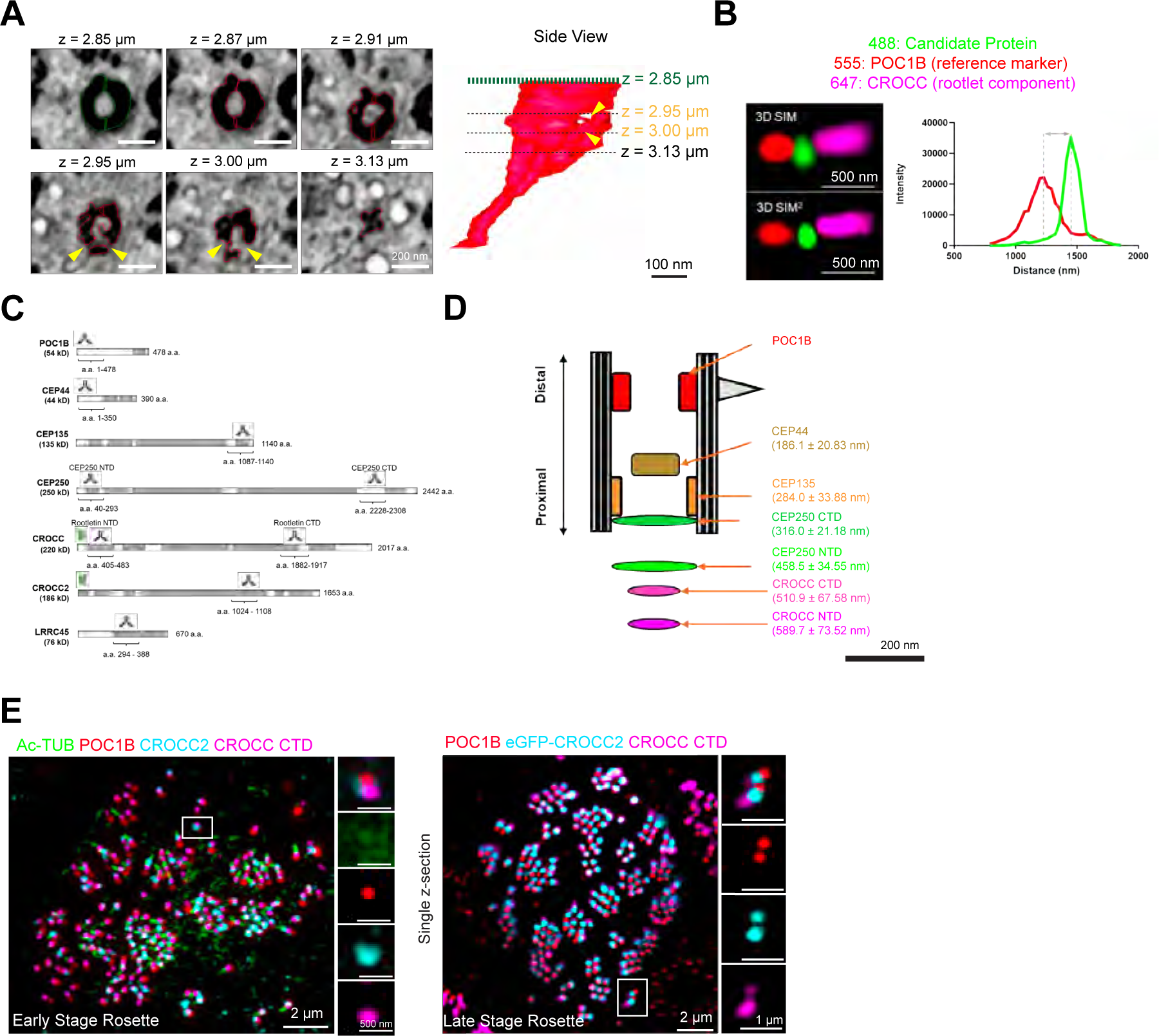
Nanoscale mapping of proteins linking basal bodies and rootlets. (A) 3D volume segmentation of rootlets emerging from the basal body. Left: 2D FIB-SEM slices sequence showing the progressive formation of the rootlet in end on view indicated by yellow arrowheads. Right: 3D volume segmentation of a rootlet as it extends toward the cytoplasm (B) Nanoscale mapping strategy using lattice 3D-SIM microscopy. The distances were measured between the fluorescence maxima of centriolar marker POC1B used as reference (555) and various markers (488) on the same z-plane. (C) Linear polypeptide sequences maps of proteins suggested forming the linker between basal bodies and rootlets. Note the regions recognised by antibodies or tagged by eGFP used in super-resolution imaging experiments. (D) Summary schematic of the relative distances of centriolar and rootlet markers to POC1B. (E) 3D-SIM micrographs of partially differentiated MCCs at the early and late-stage rosette labelled with anti-POC1B (red), eGFP-CROCC2 or anti-CROCC2 antibodies (cyan) and anti-CROCC (magenta) antibodies.

**Fig. S12.**
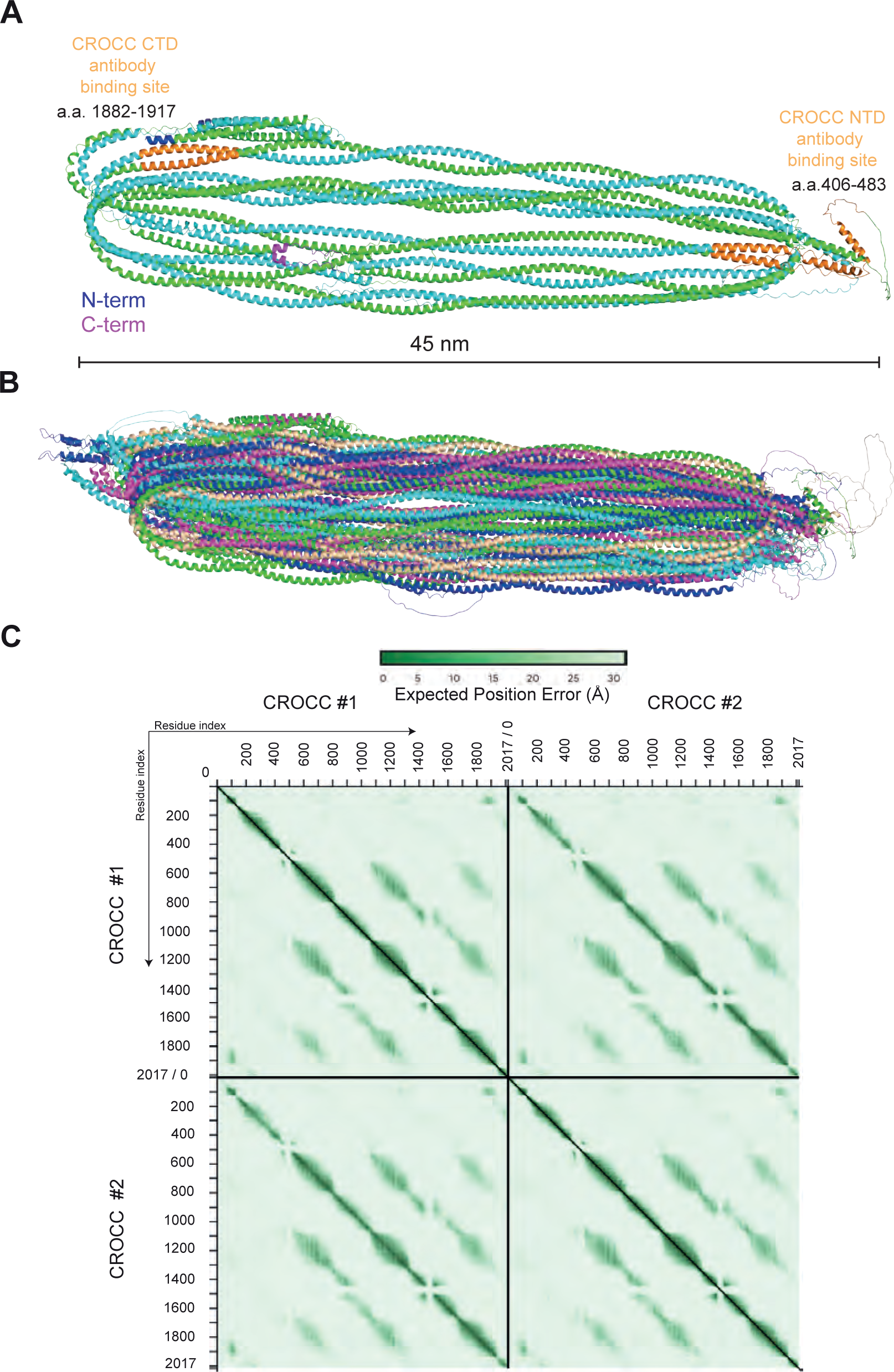
AlphaFold structure simulations of CROCC. **(A)** Cartoon representation of CROCC homo-dimer model as predicted by AlphaFold v3 with each CROCC monomer shown in a different colour. The N- and C-termini are color-coded in blue and magenta, respectively, and the length of the complex is indicated. Positions of antibody binding sites in N-terminal and C-terminal domains of CROCC are shown in orange. **(B)** Structural superpositioning of the 5 different models of CROCC homodimers predicted by AlphaFold v3 with each homodimer shown in a different colour. All five models reveal a folded back architecture of similar dimensions. **(C)** PAE plots show the predicted homo-dimeric CROCC complex and the interacting residues. The PAE plot assesses the confidence in the relative position of the protein residues within the predicted complex. The Y- and the X-axes show the residues indexed of the corresponding proteins as indicated. The aligned error in angstroms is color-coded according to the bar above the plot. Green colour indicates low PAE (high confidence in relative positions) and white colour indicates high PAE (low confidence in relative positions).

**Fig. S13.**
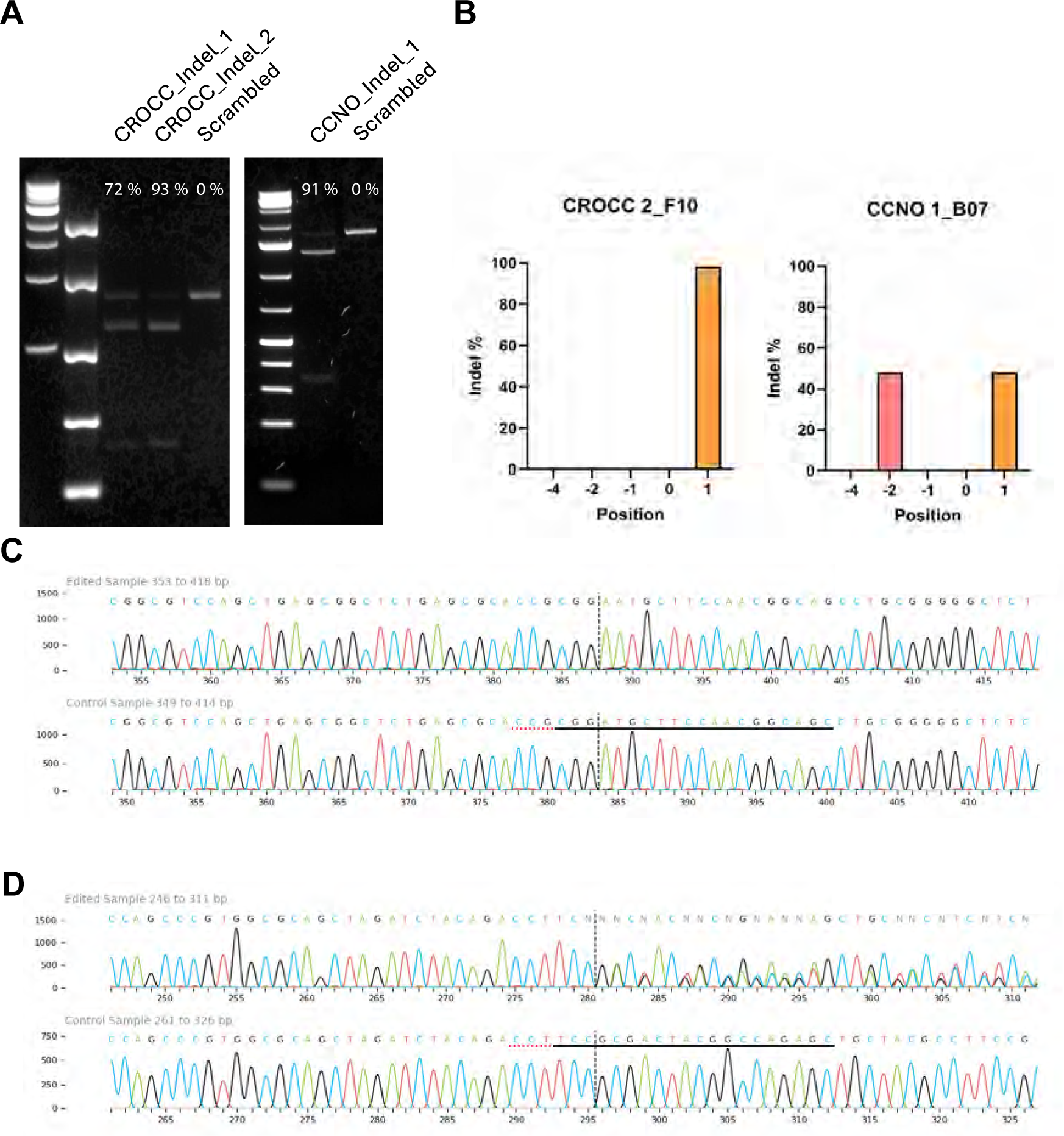
Validation of CROCC and CCNO KOs cell line by CRISPR Cas9. **(A)** sgRNA *in vitro* cleavage assay to determine cutting efficiency of selected guides to their target loci alongside a scrambled control. Efficiency is determined by measuring percentage in-lane band intensity through gel-based densitometry **(B)** Distribution of indel positions in clones derived by limiting dilution from CROCC and CCNO CRISPR-Cas9 knockout pools. **(C)** Sanger sequencing trace of CROCC 2_F10 clone highlighting target cut site via ICE analysis (Synthego). **(D)** Sanger sequencing trace of CCNO 1_B07 clone highlighting target cut site via ICE analysis (Synthego).

**Fig. S14.**
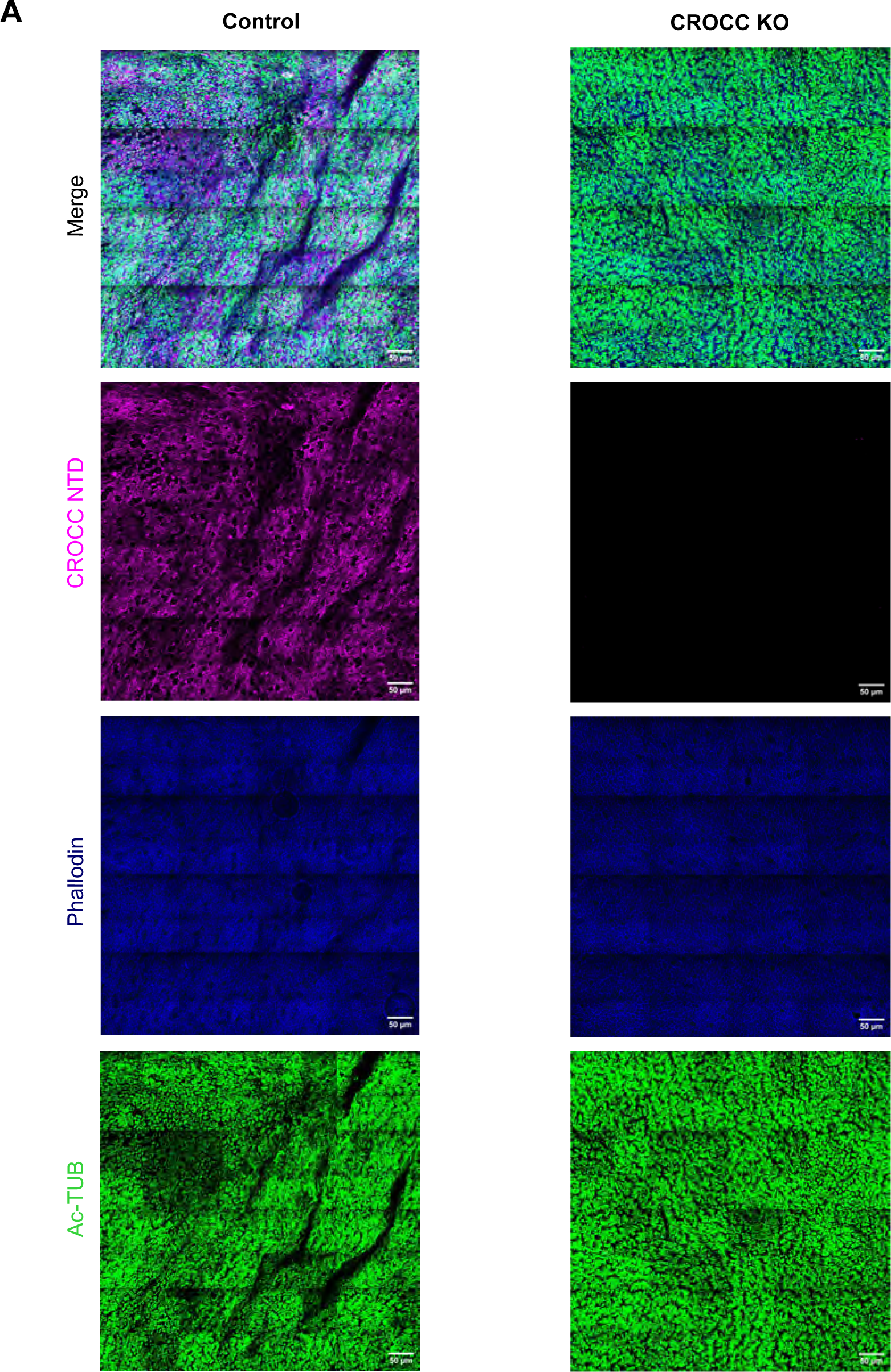
CROCC KO cells show similar ciliation level to control cells. **(A**) 3D-SIM volume maximum projection tile scan of control and CROCC KO differentiated cells labelled with anti-Ac-Tub (green), CROCC NTD (magenta), and Phalloidin F-Actin (blue).

**Fig. S15.**
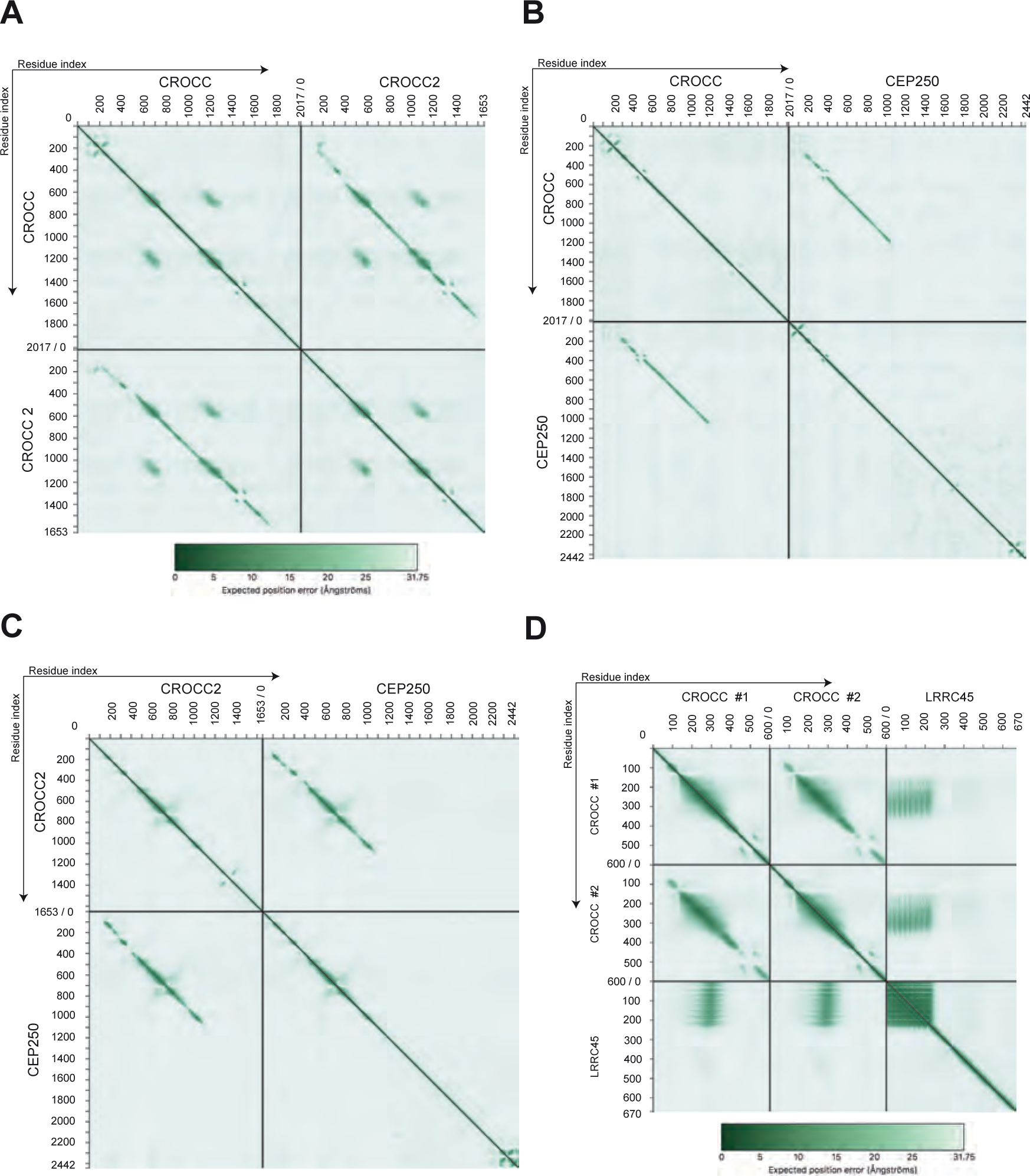
AlphaFold structure simulations of CROCC interactions with CEP250, CROCC2, LRRC45. Plots show the confidence in the relative position of the proteins within the predicted complexes. The Y- and X-axes show the residues indexed of the corresponding proteins as indicated. The aligned error in angstroms is color-coded according to the bar under the plot. Green colour indicates a low PAE score (high confidence in relative positions), and white colour indicates a high PAE score (low confidence in relative positions). In A-B, CROCC was observed to interact with CROCC2 and CEP250 and form parallel coiled-coil structures. The parallel coiled coils can be observed in the upper right and bottom left boxes. In C, CROCC2 was predicted to interact with the N-termini of CEP250 and form a parallel coiled coil, which can be observed in the upper right and bottom left boxes. In D, LRRC45 strongly interacted with the N-terminal of the dimeric coiled-coil CROCC, which can be observed in the middle left, middle right, upper right, and bottom left boxes.

**Table S1.**

Antibody list

**Movie S1-6**

**Movie S1:**

Movie of 3D volume overlaid with segmentation contours of the human airway epithelium shown in Fig.1b. Movie S1 link (DOI: <u>10.25378/janelia.26239736</u>)

**Movie S2:**

Movie of 3D volume overlaid with segmentation contours of a human airway basal stem cell shown in Fig. 1b. Movie S2 link (DOI: <u>10.25378/janelia.26239673</u>)

**Movie S3:**

Movie of segmented branching rootlet from Fig. S11h.

**Movie S4:**

Movie of 3D-SIM volume projection labelled with POC1B (cyan) and CROCC (magenta) depicting presence of free rootlets from Fig. 3g.

**Movie S5:**

Movie of 3D-SIM volume projection labelled with POC1B (cyan) and CROCC (magenta) depicting presence of free rootlets from Fig. S10b.

**Movie S6:**

Movie 3D-SIM^2^ processed eGFP-CROCC differentiated multiciliated cells.

## Notes

### Competing Interest Statement

The authors have declared no competing interest.

## References and Notes

1. Davis, J.D. and T.P. Wypych, Cellular and functional heterogeneity of the airway epithelium. Mucosal Immunol, 2021. 14(5): p. 978–990.

2. Hewitt, R.J. and C.M. Lloyd, Regulation of immune responses by the airway epithelial cell landscape. Nat Rev Immunol, 2021. 21(6): p. 347–362.

3. Spassky, N. and A. Meunier, The development and functions of multiciliated epithelia. Nat Rev Mol Cell Biol, 2017. 18(7): p. 423–436.

4. Wallmeier, J., et al., Motile ciliopathies. Nat Rev Dis Primers, 2020. 6(1): p. 77.

5. Heijink, I.H., et al., Epithelial cell dysfunction, a major driver of asthma development. Allergy, 2020. 75(8): p. 1902–1917.

6. Thomas, B., et al., Ciliary dysfunction and ultrastructural abnormalities are features of severe asthma. J Allergy Clin Immunol, 2010. 126(4): p. 722–729 e2.

7. Petit, L.M.G., et al., Airway ciliated cells in adult lung homeostasis and COPD. Eur Respir Rev, 2023. 32(170).

8. Thomas, B., et al., Dysfunctional Bronchial Cilia Are a Feature of Chronic Obstructive Pulmonary Disease (COPD). COPD, 2021. 18(6): p. 657–663.

9. Ruiz Garcia, S., et al., Novel dynamics of human mucociliary differentiation revealed by single-cell RNA sequencing of nasal epithelial cultures. Development, 2019. 146(20).

10. Herawati, E., et al., Multiciliated cell basal bodies align in stereotypical patterns coordinated by the apical cytoskeleton. J Cell Biol, 2016. 214(5): p. 571–86.

11. Lyu, Q., et al., Formation and function of multiciliated cells. J Cell Biol, 2024. 223(1).

12. Peddie, C.J., et al., Volume electron microscopy. Nat Rev Methods Primers, 2022. 2: p. 51.

13. Prescott, R.A., et al., A comparative study of in vitro air-liquid interface culture models of the human airway epithelium evaluating cellular heterogeneity and gene expression at single cell resolution. Respir Res, 2023. 24(1): p. 213.

14. Conrad, R. and K. Narayan, Instance segmentation of mitochondria in electron microscopy images with a generalist deep learning model trained on a diverse dataset. Cell Syst, 2023. 14(1): p. 58–71 e5.

15. Vijayakumaran, A., et al., Airway Cells 3D Reconstruction via Manual and Machine-Learning Aided Segmentation of Volume EM Datasets. Methods Mol Biol, 2024. 2725: p. 131–146.

16. Sorokin, S.P., Reconstructions of centriole formation and ciliogenesis in mammalian lungs. J Cell Sci, 1968. 3(2): p. 207–30.

17. Kalukula, Y., et al., Mechanics and functional consequences of nuclear deformations. Nat Rev Mol Cell Biol, 2022. 23(9): p. 583–602.

18. Li, W., et al., Nuclear localization of mitochondrial TCA cycle enzymes modulates pluripotency via histone acetylation. Nat Commun, 2022. 13(1): p. 7414.

19. Crotta, S., et al., Repair of airway epithelia requires metabolic rewiring towards fatty acid oxidation. Nat Commun, 2023. 14(1): p. 721.

20. Kunimoto, K., et al., Coordinated ciliary beating requires Odf2-mediated polarization of basal bodies via basal feet. Cell, 2012. 148(1-2): p. 189–200.

21. Nguyen, Q.P.H., et al., Comparative Super-Resolution Mapping of Basal Feet Reveals a Modular but Distinct Architecture in Primary and Motile Cilia. Dev Cell, 2020. 55(2): p. 209–223 e7.

22. Lu, Q., et al., Early steps in primary cilium assembly require EHD1/EHD3-dependent ciliary vesicle formation. Nat Cell Biol, 2015. 17(3): p. 228–240.

23. Le Guennec, M., et al., A helical inner scaffold provides a structural basis for centriole cohesion. Sci Adv, 2020. 6(7): p. eaaz4137.

24. Jain, A., et al., Mitochondrial uncoupling proteins protect human airway epithelial ciliated cells from oxidative damage. Proc Natl Acad Sci U S A, 2024. 121(10): p. e2318771121.

25. Hayes, M.J., et al., The 3D organisation of mitochondria in primate photoreceptors. Sci Rep, 2021. 11(1): p. 18863.

26. Olsson, R., The relationship between ciliary rootlets and other cell structures. J Cell Biol, 1962. 15(3): p. 596–9.

27. Vlijm, R., et al., STED nanoscopy of the centrosome linker reveals a CEP68-organized, periodic rootletin network anchored to a C-Nap1 ring at centrioles. Proc Natl Acad Sci U S A, 2018. 115(10): p. E2246–E2253.

28. Potter, C., et al., Multiple Isoforms of Nesprin1 Are Integral Components of Ciliary Rootlets. Curr Biol, 2017. 27(13): p. 2014–2022 e6.

29. Basquin, C., et al., Emergence of a Bilaterally Symmetric Pattern from Chiral Components in the Planarian Epidermis. Dev Cell, 2019. 51(4): p. 516–525 e5.

30. Mahen, R., The structure and function of centriolar rootlets. J Cell Sci, 2021. 134(16).

31. Soh, A.W.J., et al., Ciliary force-responsive striated fibers promote basal body connections and cortical interactions. J Cell Biol, 2020. 219(1).

32. Liu, Z., et al., A quantitative super-resolution imaging toolbox for diagnosis of motile ciliopathies. Sci Transl Med, 2020. 12(535).

33. Bahe, S., et al., Rootletin forms centriole-associated filaments and functions in centrosome cohesion. J Cell Biol, 2005. 171(1): p. 27–33.

34. Bond, C., et al., Technological advances in super-resolution microscopy to study cellular processes. Mol Cell, 2022. 82(2): p. 315–332.

35. Zhao, H., et al., Fibrogranular materials function as organizers to ensure the fidelity of multiciliary assembly. Nat Commun, 2021. 12(1): p. 1273.

36. Liu, Z., et al., Super-Resolution Microscopy and FIB-SEM Imaging Reveal Parental Centriole-Derived, Hybrid Cilium in Mammalian Multiciliated Cells. Dev Cell, 2020. 55(2): p. 224–236 e6.

37. Wallmeier, J., et al., Mutations in CCNO result in congenital mucociliary clearance disorder with reduced generation of multiple motile cilia. Nat Genet, 2014. 46(6): p. 646–51.

38. Chen, J.V., et al., Rootletin organizes the ciliary rootlet to achieve neuron sensory function in Drosophila. J Cell Biol, 2015. 211(2): p. 435–53.

39. Hardy, T., et al., Multisite phosphorylation of C-Nap1 releases it from Cep135 to trigger centrosome disjunction. J Cell Sci, 2014. 127(Pt 11): p. 2493–506.

40. He, R., et al., LRRC45 is a centrosome linker component required for centrosome cohesion. Cell Rep, 2013. 4(6): p. 1100–7.

41. Hossain, D., et al., Cep44 functions in centrosome cohesion by stabilizing rootletin. J Cell Sci, 2020. 133(4).

42. Ko, D., et al., Identification of a Structurally Dynamic Domain for Oligomer Formation in Rootletin. J Mol Biol, 2020. 432(13): p. 3915–3932.

43. Yang, J., M. Adamian, and T. Li, Rootletin interacts with C-Nap1 and may function as a physical linker between the pair of centrioles/basal bodies in cells. Mol Biol Cell, 2006. 17(2): p. 1033–40.

44. Kurtulmus, B., et al. LRRC45 contributes to early steps of axoneme extension. J Cell Sci 131 (2018). 10.1242/jcs.223594

45. Yang, J., et al., Rootletin, a novel coiled-coil protein, is a structural component of the ciliary rootlet. J Cell Biol, 2002. 159(3): p. 431–40.

46. Gilbert, M.C., et al., Ciliary Rootlet Coiled-Coil 2 (crocc2) Is Associated with Evolutionary Divergence and Plasticity of Cichlid Jaw Shape. Mol Biol Evol, 2021. 38(8): p. 3078–3092.

47. Carter, C.v.H.a.A.P., A cryo-ET study of ciliary rootlet organization. Elife, 2023.

48. Yang, J., et al., The ciliary rootlet maintains long-term stability of sensory cilia. Mol Cell Biol, 2005. 25(10): p. 4129–37.

49. Löschberger Anna, N.Y., Netz Ralf, Spindler Marie-Christine, Benavente Ricardo, Klein Teresa, Sauer Markus, Dr. Kleppe Ingo, Super-Resolution Imaging by Dual Iterative Structured Illumination Microscopy. 2021.

50. Galati, D.F., et al., DisAp-dependent striated fiber elongation is required to organize ciliary arrays. J Cell Biol, 2014. 207(6): p. 705–15.

51. Junker, A.D., et al., Basal bodies bend in response to ciliary forces. Mol Biol Cell, 2022. 33(14): p. ar146.

52. Werner, M.E., et al., Actin and microtubules drive differential aspects of planar cell polarity in multiciliated cells. J Cell Biol, 2011. 195(1): p. 19–26.

53. Coulon, J., J.P. Arsanto, and Y. Thouveny, Striated ciliary root-golgi association in branchial crown epithelial cells ofOwenia. Visualization of Ca2+-binding sites and ATPase activities. Protoplasma, 1986. 130(2): p. 108–119.

54. Su, E., C. Villard, and J.B. Manneville, Mitochondria: At the crossroads between mechanobiology and cell metabolism. Biol Cell, 2023. 115(9): p. e2300010.

55. Walters, M.S., et al., Generation of a human airway epithelium derived basal cell line with multipotent differentiation capacity. Respir Res, 2013. 14(1): p. 135.

56. Munye, M.M., et al., BMI-1 extends proliferative potential of human bronchial epithelial cells while retaining their mucociliary differentiation capacity. Am J Physiol Lung Cell Mol Physiol, 2017. 312(2): p. L258–L267.

57. Concordet, J.P. and M. Haeussler, CRISPOR: intuitive guide selection for CRISPR/Cas9 genome editing experiments and screens. Nucleic Acids Res, 2018. 46(W1): p. W242–W245.

58. Morone, N., et al., A Novel Sandwich Method for Serial Block Face SEM Imaging of Airway Multiciliated Epithelium. Methods Mol Biol, 2024. 2725: p. 121–129.

59. Peretti, D., et al., RBM3 mediates structural plasticity and protective effects of cooling in neurodegeneration. Nature, 2015. 518(7538): p. 236–9.

60. Kremer, J.R., D.N. Mastronarde, and J.R. McIntosh, Computer visualization of three-dimensional image data using IMOD. J Struct Biol, 1996. 116(1): p. 71–6.

61. Vijayakumaran, A., et al., Airway Cells 3D Reconstruction via Manual and Machine-Learning Aided Segmentation of Volume EM Datasets. Methods Mol Biol, 2024. 2725: p. 131–146.

62. Sikkema, L., et al., An integrated cell atlas of the lung in health and disease. Nat Med, 2023. 29(6): p. 1563–1577.

63. Ollion, J., et al., TANGO: a generic tool for high-throughput 3D image analysis for studying nuclear organization. Bioinformatics, 2013. 29(14): p. 1840–1.

64. Liu, Z., et al., A quantitative super-resolution imaging toolbox for diagnosis of motile ciliopathies. Sci Transl Med, 2020. 12(535).

65. Jeong, I., Hansen, J. N., Wachten, D. & Jurisch-Yaksi, N. Measurement of ciliary beating and fluid flow in the zebrafish adult telencephalon. STAR Protoc 3, 101542 (2022). 10.1016/j.xpro.2022.101542

66. Abramson, J., et al., Accurate structure prediction of biomolecular interactions with AlphaFold 3. Nature, 2024.

